# Gliding in the Amazonian canopy: adaptive evolution of flight in *Morpho* butterflies

**DOI:** 10.1101/2021.03.22.436469

**Authors:** Camille Le Roy, Dario Amadori, Samuel Charberet, Jaap Windt, Florian T. Muijres, Violaine Llaurens, Vincent Debat

## Abstract

The diversity of flying animals suggests that countless combinations of morphologies and behaviors have evolved with specific lifestyles, thereby exploiting diverse aerodynamic mechanisms. Elucidating how morphology, flight behavior and aerodynamic properties together diversify with contrasted ecologies remains however seldom accomplished. Here, we studied the adaptive co-divergence in wing shape, flight behavior and aerodynamic efficiency among *Morpho* butterflies living in different forest strata, by combining high-speed videography in the field with morphometric analyses and aerodynamic modelling. By comparing canopy and understory species, we show that adaptation to an open canopy environment resulted in increased glide efficiency. Moreover, this enhanced glide efficiency was achieved by different canopy species through strikingly distinct combinations of flight behavior, wing shape and aerodynamic mechanisms, highlighting the multiple pathways of adaptive evolution.

**One Sentence Summary:** By combining high-speed videography, geometric morphometrics and computational aerodynamic modelling, our study of wild Amazonian *Morpho* butterflies reveals a strong contrast between the efficient gliding flight of canopy species and the powerful flapping flight of understory species, pointing at a combined adaptive divergence of wing shape and flight behavior among sympatric species flying in different forest strata.

## Main Text

Insects display a large diversity of flight patterns reflecting their highly diversified ecologies, from the sustained, energy-efficient flight of long-range migrating species, like the desert locust (*1*) or the monarch butterfly (*2*), to the highly-manoeuvrable hovering of nectar-feeding moths (*3*), bees (*4*) and flies (*5*). This large diversity of flight modes stems from different morphological and behavioral adaptations, improving different flight performance metrics such as speed, manoeuvrability, take-off or energetic efficiency. Investigating insect flight aerodynamics is therefore crucial to understand how natural selection shapes the diversity of flight. Although insect flight has been studied in detail in several, highly divergent species, including *Drosophila* (*6*), mosquitoes (*7*) and hawkmoths (*8*), only the comparison of closely-related species adapted to different habitats can unravel the impact of ecological constraints on the diversification of aerodynamic properties.

Here, we addressed the ecological, behavioral and morphological bases of the diversification of flap-gliding flight in closely-related butterfly species. Butterflies are the only insects that regularly use flap-gliding flight, by combining periods of flapping flight interspersed with glide phases (*9*). In contrast, many intermediate-sized bird species use flap-gliding flight to reduce energetic expenditure when the aerodynamic efficiency of gliding phases is high enough (*10*). In this study, we assessed the diversity of flap-gliding flight in the neotropical butterfly genus *Morpho*. Sympatric *Morpho* species display remarkably contrasted ecologies, some species flying in the dense vegetation of the understory and others in the open canopy (*11*). Contrasted selective pressures among microhabitats may trigger divergence in flight behavior among species. Open habitats may favor a more extensive gliding behavior in species living in the canopy (*11*). We tested the divergence of flight behavior among habitats and asked whether the evolution of gliding flight in canopy species was enabled by an increased aerodynamic efficiency through changes in wing shape (*12*).

We performed a series of field and semi-field experiments in Amazonian Peru. Here, up-to-twelve *Morpho* species co-occur, allowing to investigate how specialization into different habitats impacts the evolution of flight in closely-related species in sympatry. First, we used high-speed videography to track and characterize flight behavior of wild individuals in the field and in a large insectary. We then performed geometric morphometrics analyses to precisely quantify the wing shape of these filmed butterflies and assess its covariation with flight. Finally, we used computational fluid dynamics (CFD) modelling to assess the aerodynamic efficiency associated with the contrasted wing shapes of species specialized in different habitats.

Using high-speed videography, we recorded 136 sequences of 80 wild *Morpho* butterflies freely patrolling in nature, including four understory and three canopy species (Fig. 1). From the temporal positions of each wing stroke, we measured the flapping frequency *f*, the duration of gliding phases *T*_glide_, and the temporal flap-gliding ratio *R*_glide_ of each flight (Fig. 1). While little variation was found in flapping frequency, two of the canopy species studied (*M. cisseis* and *M. telemachus*) showed sharply longer gliding phases than all understory species, spending about half of their time gliding. The third canopy species, *M. rhetenor*, strikingly differed from the other canopy species, showing surprisingly limited use of gliding, even less than all understory species (Fig. 1). These findings corroborate field observations (*11*) and highlight that contrasted flight behaviors exist within the canopy clade.

**Fig. 1.**
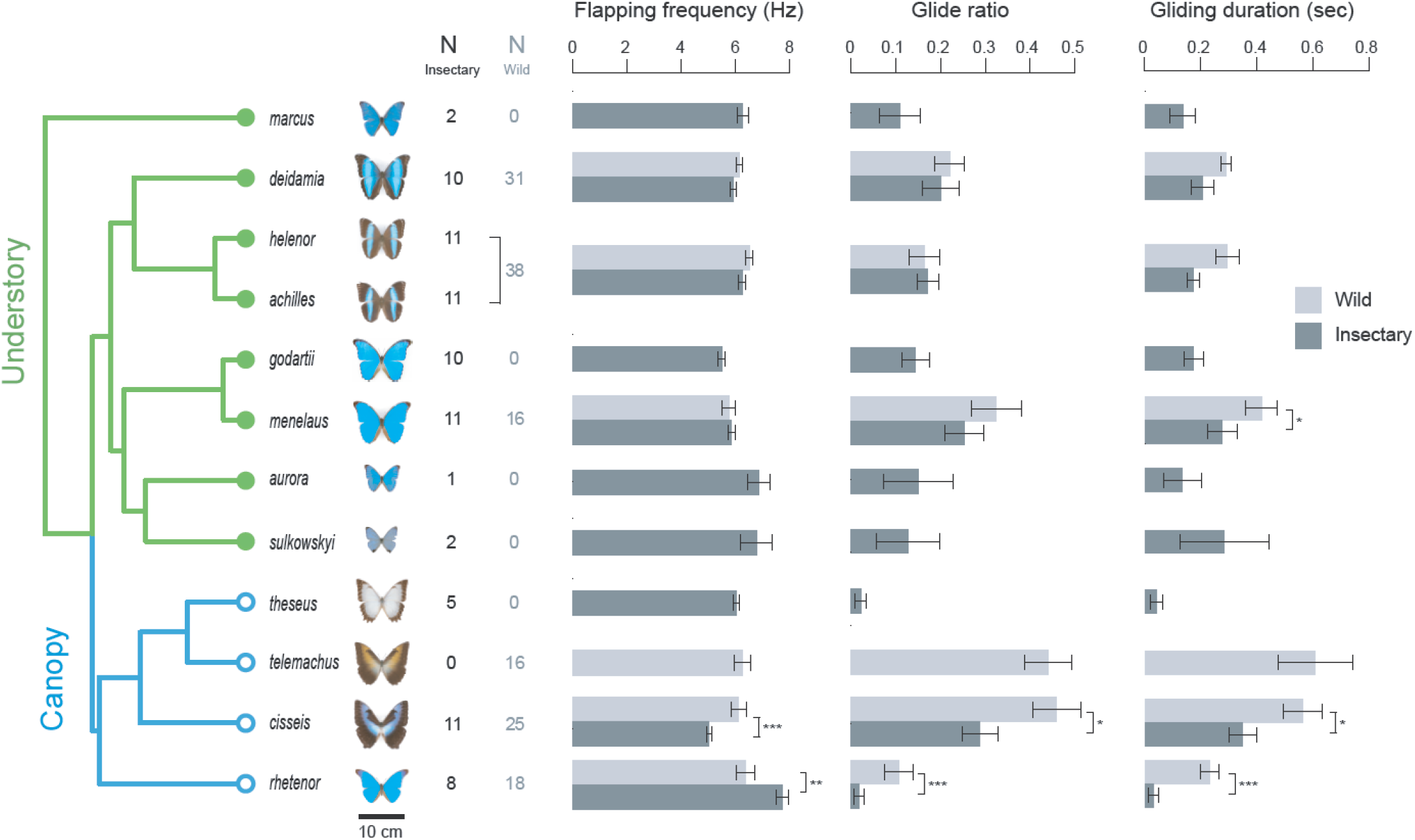
Butterflies from *Morpho* species living in the canopy use flap-gliding flight in a larger extent than species living in the understory. The twelve *Morpho* species studied here (out of 30 in the genus) are represented with their phylogenetic relationships. Differences in flap-gliding parameters measured on butterflies from different microhabitat were more striking in nature, as captivity reduced the extent of glide in canopy species. Bars indicate the mean ± standard error for each parameter, and stars indicate significant difference between nature and captivity. Note that *M. helenor* and *M. achilles* cannot be distinguished during flight: corresponding data in nature were thus pooled.

To finely characterize flight behaviors of canopy and understory species, we built a large outdoor insectary equipped with a high-speed stereoscopic videography system. In this large cage, we tracked the three-dimensional kinematics of 241 flights from 82 wild-caught *Morpho* butterflies from eight understory and three canopy species (Fig. 2A, B). We then characterized all flights using eleven flight kinematics parameters, of which six characterized the complete flap-gliding flight, three the flapping-phase and two to the gliding phase (Supplementary Methods and Table S1).

**Fig. 2.**
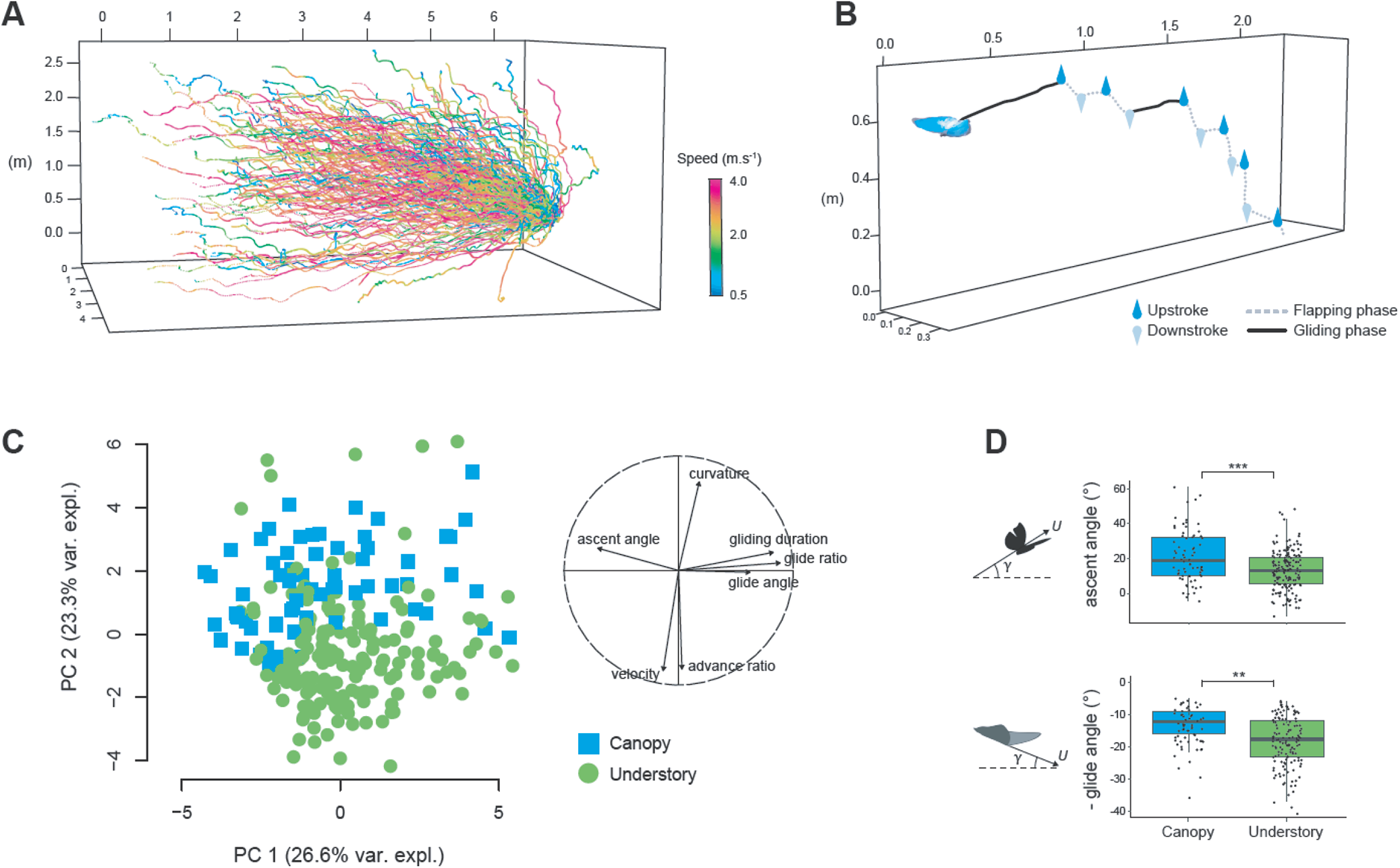
Three-dimensional flight kinematics revealed that canopy and understory species differ in various aspects of flight behavior and performance. (**A**) The 241 trajectories analyzed are shown ogether. Color indicates instantaneous flight speed. (**B**) A single flight trajectory (duration = 1.71 sec). Droplets indicate the uppermost and lowermost wing positions during up-stroke and downstroke, respectively. Gliding and flapping phases were distinguished based on wing stroke position along the trajectory. (**C**) Principal Component Analysis (PCA) showing the divergence of flight between canopy and understory butterflies. The contributions of flight parameters to PC axes are shown. (**D**) Gliding and climbing efficiency (measured as lower glide angle during gliding phase and higher ascent angle during flapping phases, respectively) were higher in canopy species, and found to diverge more strongly than expected from phylogenetic distance.

We first compared insectary flights to those recorded in the wild to assess the impact of captivity on flight behavior (Fig. 1, Table S2). Captivity reduced the extent of flap-gliding flight in canopy species. Understory species were surprisingly little affected, suggesting that they may be accustomed to flying in confined spaces. Overall, the interspecific variation in flight behavior in the insectary was broadly consistent with that observed in the wild (Fig. 1).

A Principal Component Analysis (PCA) performed on the eleven kinematics parameters showed that flight behaviors significantly differ between butterflies from the canopy and the understory (Fig. 2C). A phylogenetic MANOVA confirmed that this difference is higher than expected from a Brownian model of character evolution (Table S3). This strong divergence in flight mode between canopy and understory species therefore cannot be explained by their phylogenetic divergence only, therefore pointing at an effect of the contrasted selection regimes acting on flight evolution in the two microhabitats.

Principal component 1 was driven by the relative use of gliding flight (variation in glide-duration, glide-angle and glide-ratio), which was comparable in both canopy and understory species when flying in captivity (Fig. 2C, PC1). Principal component 2 (Fig. 2C, PC2) reflected the aerodynamic force production during flapping flight, and clearly opposed canopy and understory species: fast flight and high advance ratio for understory species on the negative values and slow flight and curvy trajectories for canopy species on the positive values (Fig. 2C, PC2). Understory species thus exhibit a more powerful wingbeat, producing higher aerodynamic forces and leading to higher advance ratios and straighter high-speed flights.

Glide angle and ascent angle were significantly more divergent between canopy and understory species than predicted by phylogenetic distances (Fig. 2D, Table S3), suggesting a strong effect of natural selection on these two flight components. During the few flapping phases they performed, canopy butterflies also climbed more steeply: the mean ascent angle was 70% larger in canopy than in understory species (Fig. 2D; γ_ascend,canopy_=22°±3°, *n*=70 flapping phases; γ_ascend,understory_=13°±1°, *n*=171 flapping phases). Thus, although understory species tended to fly at higher advance ratio and flight speed (Fig. 2C, PC2), the ascent angle of canopy species was higher (Fig. 2C, PC1; Fig. 2D). This could stem from an increased behavioral tendency to fly up, and/or from a higher climbing efficiency generated by their morphology.

During the gliding phases, canopy butterflies had a 36% lower glide angle than the understory butterflies (Fig. 2D; γ_glide,canopy_=7°±2°, *n*=61 gliding phases; γ_glide,understory_=11°±1°, *n*=135 gliding phases). Such shallower glides allow canopy butterflies to travel longer distances for a given height loss, consistent with the longer gliding phases measured in the wild. Glide angle is directly related to the aerodynamic efficiency parameter lift-to-drag ratio (*13*), and shallower angles detected in canopy species might be promoted by their divergent wing shapes. This combination of field and semi-field experiments shows that the evolutionary shift from understory to canopy resulted in an increased use and efficiency of gliding flight (Fig. 2D), combined with a reduction in aerodynamic force production during forward flapping flight (Fig. 1C, PC2).

We then investigated the contribution of morphological divergence in the adaptive evolution of flight between canopy and understory species. We precisely quantified wing shape of the filmed butterflies using geometric morphometrics and detected a strong covariation between wing shape and flight behavior using a phylogenetic partial least square analysis (Fig. 3A). Butterflies with more rounded wings and higher wing-loading (*WL*) flew at higher flight speeds and advance-ratio, and accelerated more rapidly (Fig. 3A and Table S4). These results suggest that the evolution of smaller (high *WL*), more-rounded wings indeed increased force production during flapping flight (Fig 3A). Our analyses demonstrate that flight power therefore tightly co-evolves with wing shape. In contrast to flapping flight parameters, gliding parameters were weakly correlated with wing shape and aspect-ratio *AR* (Fig. 3B, Table S4). Canopy species are efficient gliders yet exhibiting strikingly diverse wing-loadings, aspect-ratios and wing shapes (Fig. 3): two of the studied canopy species are slow flyers with low aspect-ratio, low wing-loading triangular wings (*M. cisseis*, and *M. theseus*), whereas the fast-flying *M. rhetenor* has high aspect-ratio elongated wings with high wing-loading. This begs the question of how this divergence in wing shape among species altered gliding efficiency.

**Fig. 3.**
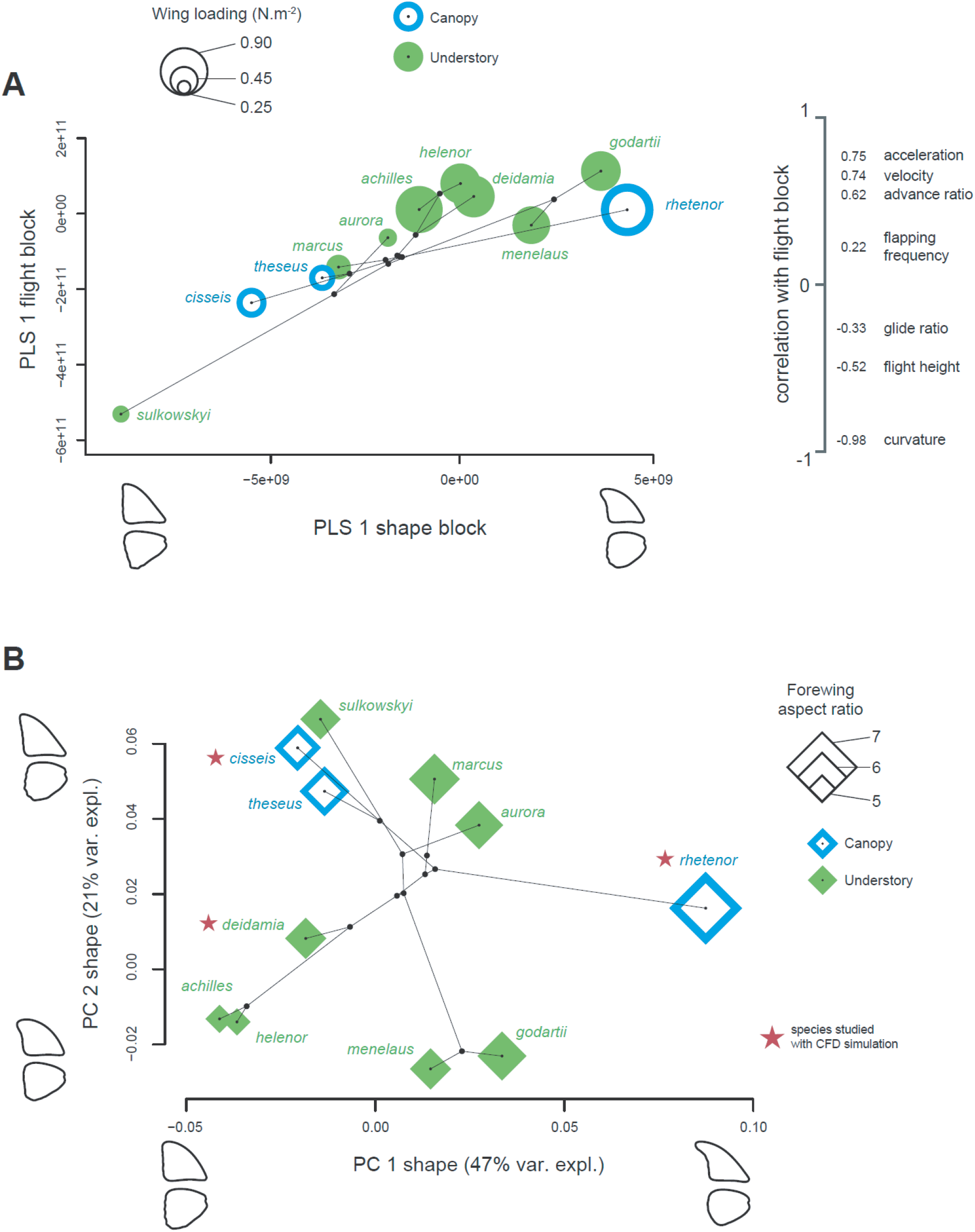
Wing shape and wing-loading jointly covary with flight behavior. (**A**) Phylogenetic partial-least square (PLS) analysis shows a tight covariation between wing shape and flight behavior (r-PLS = 0.89; P = 0.02; 74% of covariation explained). This covariation opposes triangular to rounded wings, respectively associated with slow – curvy flight, and straighter – more powerful flight. Wing-loading (depicted by symbol size) also covaries with flight, suggesting that the evolution of flight is tightly linked to both wing shape and body morphology. Flight loadings are indicated on the right. (**B**) Phylogenetic morphospace depicting variation in wing shape among species, diamond size indicates aspect ratio

To answer this, we used Computational Fluid Dynamics (CFD) to determine how gliding performances differ between three *Morpho* species with strikingly different wing shape and flight behavior (Fig. 3B): *M. cisseis*, a slow-flying canopy species with large triangular wings; *M. rhetenor*, the fast-flying canopy species with markedly elongated wings; and *M. deidamia*, a fast-flying understory species with rounded wings.

For each species, we produced *in-silico* wings based on our gliding flight experiments (Figs. 4A, S2). We then performed gliding-flight CFD simulations to determine the lift-to-drag ratio to angle-of-attack curves (*L/D-*α, Fig. 4B). For all three species, maximum lift-to-drag ratio *L/D*_max_ was attained at α=6° (Fig. 4B). Both canopy species (*M. cisseis* and *M. rhetenor*) showed ∼12% greater *L/D*_max_ than the understory species (*M. deidamia*) (*M. cisseis*: *L/D*_max_=5.83; *M. rhetenor*: *L/D*_max_=5.71; *M. deidamia*: *L/D*_max_=5.18), indicating that their wing shapes confer higher gliding efficiency.

**Fig. 4.**
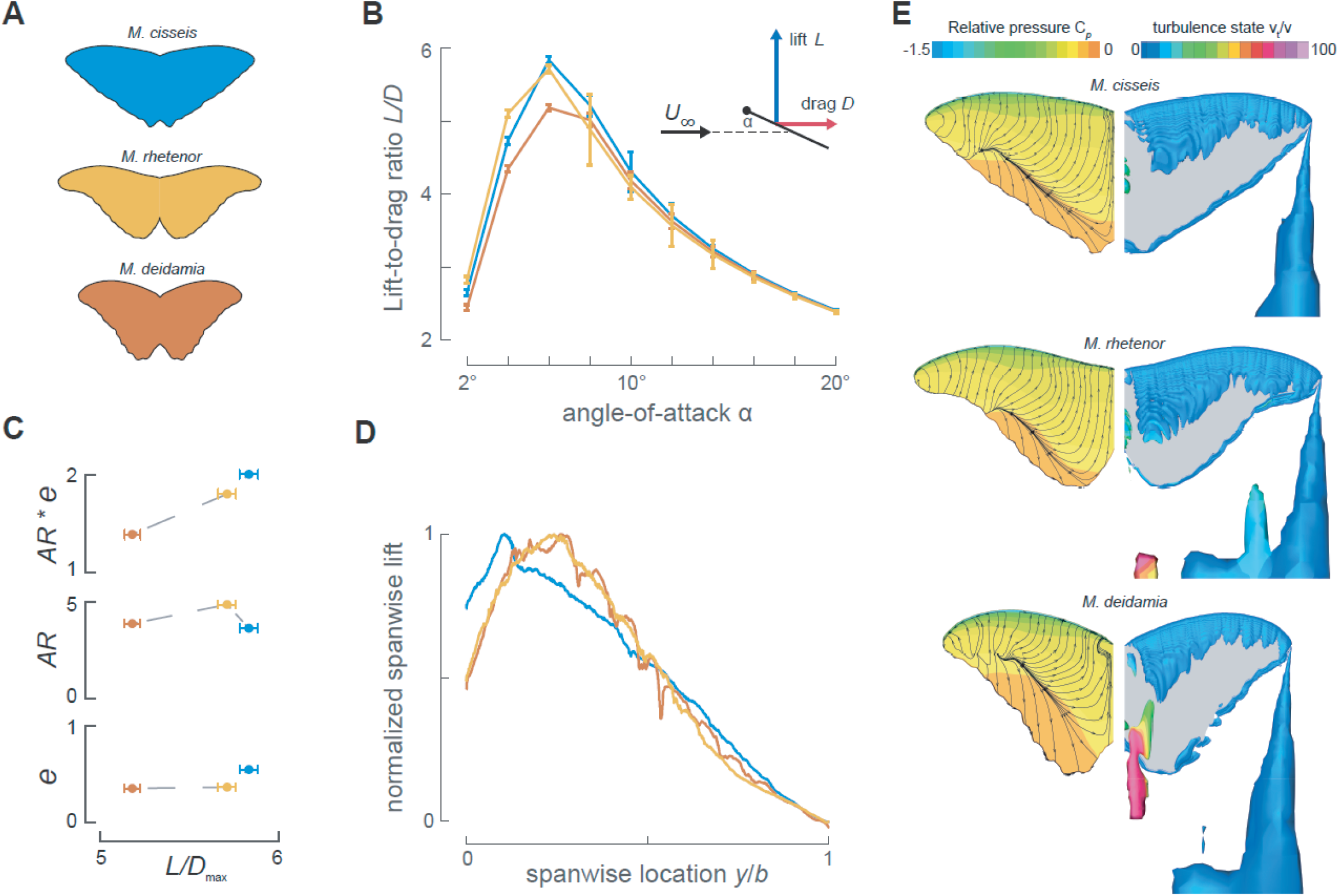
Higher gliding efficiency in canopy species is achieved through different wing shapes. (**A**) Gliding wing shape of the three species studied with Computational Fluid Dynamics (CFD). The position of the wings during gliding flight was obtained from top view filming in the insectary (Fig. S2). (**B**) Lift-to-drag ratio as a function of angle-of-attack. The two canopy species (*M. cisseis* and *M. rhetenor*) show a one-unit greater maximum lift-to-drag ratio. Error bars show numerical uncertainties obtained by dedicated verification studies (Supplementary Methods). (**C**) Aerodynamic efficiency, (**D**) spanwise lift distribution and (E) flow-fields around Morpho wings at maximum lift-to-drag ratio. (**C**) Aerodynamic efficiency equals the product of wing aspect-ratio *AR* and span-efficiency *e*. Canopy species *M. rhetenor* and *M. cisseis* have increased efficiency due to higher *AR* and *e*, respectively

Wing shape primarily affects induced drag of a wing, which inversely scales with the product of wing aspect-ratio *AR* and span-efficiency *e* (*14*). Therefore, we tested how these parameters varied between species (see Supplementary Methods).

The wing planform of *M. rhetenor* has a higher aspect-ratio than the other two species (Fig. 4A,C), whereas *M. cisseis* has a higher span-efficiency than *M. rhetenor* and *M. deidamia* (Fig. 4C). Total induced aerodynamic efficiency, as resulting from the product of span-efficiency and aspect-ratio, was higher in the two canopy species, consistent with their higher *L/D*_max_ (Fig 4C). These results provide functional evidence that wing shape divergence among *Morpho* species directly affects gliding efficiency. Interestingly, they also show that different canopy species with contrasted wing shapes achieve similarly high *L/D*_max_ through different aerodynamic mechanisms.

Airflow visualizations at *L/D*_max_ (Fig. 4E) show that all gliding butterflies produce a stable Leading-Edge-Vortex and streamwise wingtip vortices. *M. deidamia* and *M. rhetenor* produce additional streamwise vortices: a wing root vortex and a vortex at two-third span, respectively (*15*). This explains why these high-wing-loading species have such variable spanwise lift-distribution and consequently a reduced span-efficiency (Fig. 4C,D) (*15*).

Our combination of aerodynamic and ecological approaches revealed how natural selection imposed by different microhabitats can drive the evolution of flap-gliding flight in jointly altering wing shape and flight behavior. Butterflies from species evolving in the cluttered understory habitat display short gliding phases and powerful flapping phases, resulting in high flight speeds and advance ratios. In contrast, evolution in the open canopy resulted in a more efficient gliding flight, illustrated by the reduced descend angles during gliding phases observed in the canopy species.

Comparing the aerodynamic performances of closely related species adapted to different ecological conditions allowed us to identify how subtle differences in wing shape affect flight. The fact that canopy species with contrasted flight behaviors and morphologies achieve similar gliding performances through different aerodynamic mechanisms in turn illustrates how adaptive evolution is fueled by the flexible adjustment of morphology, behavior and performance.

## Acknowledgments

The authors thank the Peruvian authorities, and in particular SERFOR for providing the necessary research permits (permit: 002-2015-SERFOR-DGGSPFFS). We thank Ronald Mori Pezo and Joel Pintado for help with the butterfly sampling. We thank Tyson L. Hedrick and Guilherme Vas for their advice on the videography and CFD analyses, respectively.

## Author contributions

C.L.R., V.L., and V.D. designed the research plan and performed the field experiments. S.C. analyzed experimental field data. D.A., F.T.M., and J.W. performed the CFD study. C.L.R., V.L., V.D. and F.T.M. wrote the paper. All authors edited the manuscript.

## Materials and Methods

### Studying flight behavior, kinematics and performance in field and semi-field

Our field study was performed from July to September 2017 in the regional park of the Cordillera Escalera (San Martin Department), in the North of Peru (06°27′07″S, 76°20′47″W; ca. 450 m a.s.l.). This area is among the most *Morpho*-species-rich parts of Amazonia (*1*). We studied exclusively males, as females were too rarely seen due to their cryptic flight behavior, contrasting with the extensive patrolling displayed by males. Only undamaged specimens were selected. Wing damage significantly affect flight performance (*2, 3*) and hence would have obscured the link between wing shape and flight dynamics.

Our field work consisted of two sub-studies: In *Sub-study 1* we used high-speed videography to observe flap-gliding flight in the wild of 80 specimens belonging to four understory species (*M. helenor*, *M. achilles*, *M. deidamia*, *M. menelaus*) and three canopy species (*M. rhetenor*, *M. cisseis*, *M. telemachus*). In *Sub-study 2* we used an outdoor insectary equipped with a stereoscopic high-speed videography system (Fig. S1) to study in detail the three-dimensional kinematics of 241 flights of 82 *Morpho* butterflies, belonging to eight understory species and three canopy species.

For the semi-field study, butterflies were captured in the wild, which occurred in unbalanced proportions: about 10 individuals were captured for the most commonly encountered species, i.e. *M. helenor*, *M. achilles*, *M. menelaus*, *M. godartii*, *M. deidamia*, *M. cisseis*, *M. rhetenor*, whereas fewer individuals were captured in rarely encountered species such as *M. marcus* (n=2), and for species inhabiting the less-accessible cloud forest i.e. *M. theseus*, *M. sulkowskyi*, *M. aurora* (n=5, 2 and 1 respectively) (Fig. 1). Species inhabiting cloud forests were collected in the Alto Mayo region, near the village of El Afluente (5°39′45″S, 77°41′47″W; ca. 1300 m a.s.l.), located 230 km North-West from Tarapoto.

#### Sub-study 1. Quantifying flap-gliding flight behavior in the wild

To investigate flap-gliding behavior in natural conditions, flight sequences were recorded in the wild using a handheld high-speed camera (Gopro Hero5 Black set at 120 images per second). A total of 136 natural flight sequences were recorded (mean duration of 2.81 ± 2.53 s, ranging from 0.6 to 16.0 s.), covering three canopy and three understory species. In each video, we manually identifying the start of the downstroke and upstroke of each wingbeat. From this, we separated the flight sequence in to flapping phases and gliding phases. Gliding phases were defined as periods in which the wings were not flapping for at least 0.17 s (20 video frames). For each flapping phase, we determined the average wingbeat frequency as *f = N*_wb_ / *T*_flap_, where *N*_wb_ is the number of wingbeats in the flapping phase and *T*_flap_ is the duration of the flapping phase. For the gliding phases, we determined the gliding duration (*T*_glide_). From this, we determined glide ratio of the flap-glide flight as *R*_glide_ = Σ*T*_glide_ / *T*_total_, where *T*_total_ is the length of the digitized video.

#### Sub-study 2. Studying flap-gliding flight behavior, kinematics and performance in in semi-field conditions

To study flap-gliding flight of *Morpho* butterflies in detail, we built a large outdoor insectary (9m × 4m × 2.5m) equipped with stereoscopic high-speed videography system for recording flying butterflies (Fig. S1). The insectary was located in a sheltered spot unexposed to wind close to the field study site.

Prior to each experiment, we captured butterflies mainly at the same site as in *Sub-study 1*. Captured specimens were released from a shaded side of the insectary, and generally flew towards the sunniest part of the flight arena. We filmed the flights using the videography system, consisting of three orthogonally positioned video cameras (Gopro Hero5 Black, recording at 240 images per second). Two cameras were mounted horizontally on tripods, and provided a side and front view of the flying butterfly. The third camera was fixed to the ceiling facing downwards, and thus provided a top view (see Movie S1 and S2 for a typical flight sequences). Only trials in which butterflies flew across the insectary were used for analysis, thus assuring comparability between flight sequences. Multiple trials were performed until three suitable sequences were obtained for each individual whenever possible. A total of 241 sequences were included in the analyses, with a mean duration of 2.16 ± 0.83 s, ranging from 0.6 to 6.0 s (Fig. 2A).

Analysis of the videography data was done in MATLAB (Mathworks Inc). Stereoscopic video sequences obtained from the three cameras were synchronized with respect to a reference frame. Prior to each filming session, the camera system was calibrated with the direct linear transformation (DLT) technique (*4*) by digitizing the positions of a wand moved throughout the insectary. Wand tracking was done using *DLTdv6* (*5*), and computation of the DLT coefficients was performed using *easyWand* (*6*). Frame distortion (fisheye effect) due to wide-angle settings was also corrected in *DLTdv6*.

After spatial and temporal calibration, we also used *DLTdv6* (*5*) to digitize the three-dimensional positions of the butterfly at each video frame by manually tracking the body centroid in each camera view. In addition, the start and end of each wingbeat was identified by manually digitizing the video frames at which the wing was at the highest upstroke positions and the lowest downstroke positions, thus transcribing the spatial and temporal position of each wing stroke along the flight trajectory (Fig. 2B).

Based on these wing stroke data, we used the same method as in the field study (*sub-study 1*) to determine the flap-gliding flight parameters wingbeat frequency *f*, glide duration *T*_glide_, and flap-gliding glide ratio *R*_glide_. This allowed for a direct comparison between flight behavior in the field and in the insectary.

Butterfly positions throughout the flight trajectory were post-processed using a linear Kalman filter (*7*), providing smoothed temporal dynamics of location **X**(*t*), velocity **U**(*t*) and acceleration **A**(*t*) of the body centroid, in the world coordinate system (Fig. S1). Based on these data, we determined flight-averaged flight height *H*, flight speed *U*, and acceleration *A*. We characterized the straightness of the trajectories using two different parameters: sinuosity and curvature (*8*) (see Table S1 for formula). For the flapping phases, we quantified propulsive performance (*9*) by the advance ratio as

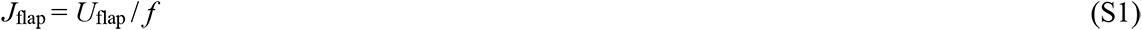

and ascent angle as

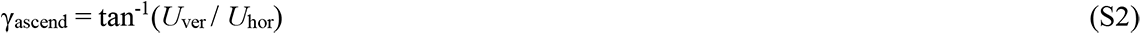

where *U*_flap_ is the average flight speed during the flapping phase, and *U*_ver_ and *U*_hor_ are the vertical and horizontal components of it, respectively. On the gliding phases, we quantified gliding efficiency using glide angle, computed as

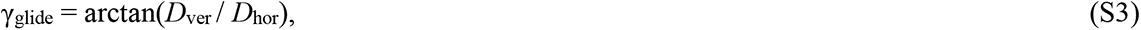

where *D*_ver_ is the vertical distance (height loss) and *D*_hor_ is the horizontal distance covered during the gliding phase.

The correlations between the measured flight parameters were all smaller than 0.7, showing that they were not fully redundant, and they were then assumed to describe at least partly different aspects of flight behavior. We retained the mean and maximum values out of the three flight sequences analyses per individual. Mean and maximum values per species were also computed to perform subsequent phylogenetic comparative analysis.

Detailed description and formula of each flight parameter measured during our experiments can be found in Table S1.

### Quantifying variation in wing shape and body morphology

After being filmed, the detached forewings and hindwings of each specimen were photographed in dorsal view using a Nikon D90 camera in controlled light conditions. We used a landmark-based geometric morphometric method (*10, 11*). Geometric morphometric has been widely used to describe biological shapes, notably insects wing (*12-14*), albeit rarely in combination with flight analysis (e.g. *15*). We described wing shape outline using 300 semi-landmarks equidistantly spaced along the wing outline. Semi-landmarks are commonly used to describe curvy shapes such as wing outlines, which lack of identifiable landmarks (*16*). We placed one fixed landmark at the wing base, fixing the overall landmark configuration, and the semi-landmarks were slid along the local tangent to the curve in an iterative process, to remove the variation along the outline due to a lack of homology (*16*). The procedure was applied to the left fore- and hindwing. All landmarks were digitized using TpsDig2 (*17*).

In order to compare wing shapes of different specimens, we then performed a Generalized Procrustes Analysis (*18*) separately on forewings and hindwings using the geomorph R package (*19*). Flight results from the joint use of the two pairs of wings, and is therefore potentially affected by their combined shape variations. To test the relationship between wing shape and flight behavior, we thus combined the two datasets as follows. After the independent superimpositions of the two sets of wings, we conducted a principal component analysis (PCA) on each set of procrustes coordinates. We then combined the principal components (PCs) of each of these two PCAs, and performed a new PCA on this global dataset to obtain PC axes combining the information of the fore- and hindwing of each individual. Wing shape variation was further described at the species level to allow phylogenetic analyses. For such analyses, we computed the mean shape of each species as their mean landmark coordinates in the individual morphospace. The between-species shape variation was described following the same process explained above, but using the mean species wing shapes instead of specimens wing shapes.

We also measured forewing aspect ratio (the ratio of wingspan to mean wing width) on each specimen using *wingImageProcessor* v.1.1 for Matlab (https://biomech.web.unc.edu/wing-image-analysis/) to investigate the relationship between flight parameters and more straightforward shape descriptor.

Wing shape is not the only morphological trait affecting flight. Body mass relatively to wing area (i.e. the wing loading) also highly constrain flight capacities (*20, 21*). Specimens bodies were thus weighted with a precision of 0.01g. We measured wing area from the digitized landmarks using *Momocs* R package (*22*) and then computed the wing loading.

### Statistical analyses

To account for the large differences of numerical range among flight parameters, the analyses described below were performed using the scaled and centered flight parameters.

#### Comparing flight behavior between nature and insectary

To assess in which extent the flight performed in insectary reliably reflects natural flight behavior, we tested the effect of captivity on the set of flight parameters measured in both conditions (i.e. wingbeat frequency, glide ratio and gliding duration) using a MANOVA, as well as specifically on each parameter using ANOVAs (Table S2). We then assessed whether understory and canopy species were similarly affected by testing the effect of captivity separately within each group.

#### Variation in flight behavior among species and microhabitats

Variation in flight behavior measured in the insectary was examined by conducting a PCA on the set of 11 parameters measured from the flight trajectories. This exploratory analysis allowed a first visual assessment of the association between microhabitat and flight behavior. Using the species mean for each flight parameters, and the latest molecular phylogeny of *Morpho* butterflies (*14*), we then tested whether closely related *Morpho* species showed more similar flight behavior than distantly related ones. This was done by computing the phylogenetic signal on each flight parameter using the Blomberg *K* (*23*), as well as on the whole set of parameters using the multivariate *K*-statistics (*24*). Phylogenetic signal was similarly computed on the flight parameters measured in the wild. We then tested the effect of microhabitat on flight behavior while controlling for the phylogenetic distance, for each flight parameter using phylogenetic ANOVAs, and all parameters together using a phylogenetic MANOVA (*25, 26*). The effect of microhabitat was tested on both insectary and wild-measured flight parameters (Table S3).

#### Effect of wing loading and aspect ratio on flight parameters

We tested the effect of wing loading and wing aspect ratio on the set of flight parameters using multivariate regressions. These effects were additionally tested while accounting for phylogeny using multivariate phylogenetic regression (function ‘procD.pgls’ of the R package *geomorph*) (*19*). In order to explore the specific effect on these morphological features on the different aspects of flight behavior, we then performed separated standard and phylogenetic linear regressions on each flight parameter (Table S4).

#### Covariation between wing shape and flight behavior

The covariation between wing shape and flight was tested accounting for the phylogenetic structure by performing a phylogenetic two-blocks partial-least squares regression (‘2B-PLS’, function ‘phylo.integration’ of the R package *geomorph* (*27*) between shape and flight data (i.e. shape PCs and flight parameters). 2B-PLS analysis focuses on the covariation between two sets of multivariate data, by constructing pairs of linear combinations of parameters within each dataset that maximally co-vary across datasets (*28*). The shape block of the PLS combines information from both the fore- and hindwing, thus preventing a direct visualization of the shape change correlated with flight. We then separately assessed the contributions of forewing and hindwing to the PLS shape axis. The corresponding shape changes were visualized by projecting the regression scores obtained from multivariate regressions of procrustes wing coordinates on the PLS shape axis (*29, 30*).

### Studying the aerodynamics of gliding flight using Computational Fluid Dynamics

Computational Fluid Dynamics (CFD) allows detailed analysis of the airflow characteristics around an object by solving the Navier-Stokes equations numerically (*31*), and has become an essential tool for the study of insect flight aerodynamics (e.g. *32, 33, 34*). We used the CFD solver ReFRESCO (https://www.refresco.org/) to study the gliding flight performance of three different *Morpho* species, in order to determine how differences in wing shape affect gliding flight performance.

To focus on the effect of wing shape on aerodynamic force production, we set all other flight characteristics to be equal between the three studied species. This includes the Reynolds number (Re), wing profile shape, wing flexibility and wing angle-of-attack. In order to keep wing profile shape and wing flexibility equal between all species, we assumed all wings to consist of a rigid flat plate. For all wings, we systematically varied the wing angle-of-attack between our simulations from α = 2° to 20° at steps of 2°. For all three species, we kept the Reynolds number (Re) constant at a value of order 5200. The Reynolds number is defined as Re = *U*_∞_ *c*/ν, with *U*_∞_ the free-stream airflow velocity, *c* the mean wing chord and ν the kinematic viscosity. Based on the mean wing chord of each wing (*c* = 4.82, 3.91 and 3.88 cm for *M. cisseis*, *M. deidamia* and *M. rhetenor*, respectively, Table S2) we set the free-stream airflow velocity for all CFD simulations as *U*_∞_ = 1.59, 1.97 and 1.98 m.s^-1^ for *M. cisseis*, *M. deidamia* and *M. rhetenor*, respectively. The methodology and supplementary results of this numerical analysis are discussed in detail below.

#### CFD model development, testing and validation

The used CFD software package ReFRESCO was developed by the Maritime Research Institute Netherlands (MARIN, Wageningen, The Netherlands) (*35*). It is in principal second order accurate using a TVD Harmonic scheme for the convective fluxes. For the convection of turbulence quantities however a first order upwind scheme was used. For the estimation of the discretization uncertainty, it is required that the iterative error is small enough such that it does not influence the discretization error: for all simulations the solution was steady, and the infinity norm dropped several orders. It was checked that a stricter convergence tolerance did not change the final uncertainty estimation. Additionally, the convergence of the computed forces was monitored. These final estimated uncertainties are depicted as error bars in the Figure 4B.

As a numerical grid, we used unstructured, hexahedral based meshes, built with Hexpress (Numeca Inc). Viscous layer grid insertion was used to be able to solve the flow down to the wall, i.e. without the use of wall functions (y^+^ < 1). Grids for different angles-of-attack of the wing were obtained using a grid deformation method. The wing (hinge-to-tip length 0.085m) was positioned in the middle of the computational domain that consisted of a rectangular box. Only one wing was modelled, exploiting symmetry. The boundary conditions were set as follows: we imposed the free-stream velocity *U*_∞_ at the inlet, no-slip condition at the wing, symmetry at the symmetry plane and zero pressure at the remaining boundaries.

We verified our CFD method, including the turbulence models and grid quality, by estimating the numerical / discretization uncertainty using a grid refinement exercise (Table S5) (*36*). First, we checked that the boundaries were sufficiently far away from the wing: increasing the domain with a factor 1.5 decreased the drag and lift by less than 0.2%. Then, we systematically increased the number of cells in the numerical grid using our grid refinement exercise (Table S5). For the model selection we did this in four steps, and for the final study we used five steps per species (Table S5). Our analysis showed that the coarsest grids were already fine enough to capture the geometry of the wing and to resolve the flow around the wing sufficient accurately (Fig S2D). We used the simulation with the finest grid to determine aerodynamic forces and corresponding airflow dynamics; we estimated the uncertainties in aerodynamic force production using the complete set of simulations at all grid sizes (*36*).

All CFD data were described in the world reference frame, with the *x*-axis in the streamwise direction, the *y*-axis along the wingspan and the *z*-axis vertically up. Velocities are defined in this reference frame as **U** = [*u*, *v*, *w*], and aerodynamic forces on the wing as **F**_aero_ = [*F_x_*, *F_y_*, *F_z_*].

#### Turbulence model selection analysis

At the Reynolds number regime of order 5000 at which Morpho butterflies glide, the flow around the wing changes from laminar to turbulent and therefore aerodynamic performance is notoriously difficult to quantify (*37*). Therefore, selection of an appropriate turbulence model needs to be done carefully. To do so, we tested four turbulence model configurations; for each model the numerical uncertainty was determined using a dedicated grid density study. The final turbulence model selection was done by comparing the lift-to-drag ratio from each simulation with that of a real butterfly estimated from the semi-field experiments.

For this, we focused specifically on the glide sequence of *M. cisseis* from which also the gliding-flight wing shape was determined (Fig. S2). From the wing orientation and glide velocity we determined the wing dihedral angle as Γ = 16° and angle-of-attack as α = 11°. Using Newton’s second law of motion, we determined the aerodynamic force vector during the gliding flight as

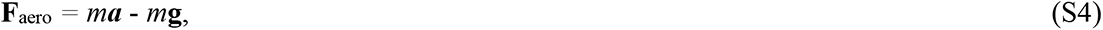

Where **a** is the body acceleration vector and **g** is the gravitational acceleration vector. We then estimated the lift force *L* and drag force *D* as the aerodynamic force components normal and tangential to the velocity vector (Fig. S1B). The lift produced by single wing was then estimated as

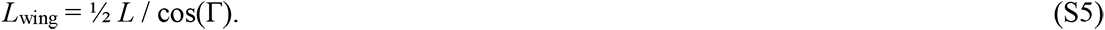

Based on these, we determined the lift coefficient and drag coefficient as

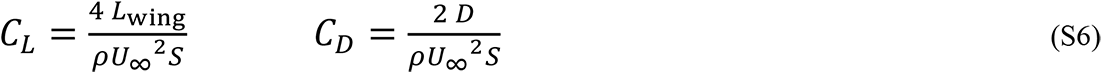

where ρ is air density and *S* is wing surface area. The lift-to-drag ratio was estimated as

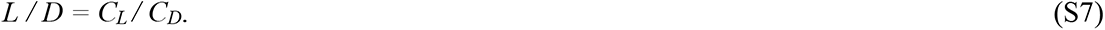

We then performed CFD simulations of this *M. cisseis* wing shape operating at the same flight speed and angle-of-attack, with various turbulence models. For each turbulence model configuration, we determined the lift coefficient, drag coefficient and lift-to-drag ratio from the CFD simulation. The uncertainty interval for the aerodynamic forces and corresponding coefficients estimated in our CFD were determined using on a grid refinement exercise (*36*). By comparing the forces from the different models with the empirical estimate, we determined which model was best.

#### Selection of the best turbulence model

We tested two turbulence models in four configurations. The turbulence models were the SST k-ω 2003 turbulence model and the Spalart-Allmaras model (*38*). We chose these models because they were both developed for aerodynamic purposes specifically and because they have been extensively validated in literature. Both models do not solve transition explicitly, and therefore we also used the SST k-ω together with the *γ*-Re_θ_ equations for transition, to investigate whether transition plays an important role in the definition of the flow around the wing. Given the sensibility of the latter model to inflow conditions, we considered two different setups, with inlet values of *ν_t_*/*ν* of 2.5 and 10, following experimental results in (*39*). In summary, we thus tested the following four turbulence model configurations:

1. The Spalart-Allmaras model
2. The SST *k*-ω model without transition model
3. The SST *k*-ω with *γ*-Re_θ_ transition model and *ν_t_* / *ν* = 2.5 inflow condition
4. The SST *k*-ω with *γ*-Re_θ_ transition model and *ν_t_* / *ν* = 10.0 inflow condition.

All initial calculations converged to a steady state. Figure S3 shows the relative pressure and limiting streamlines on the wing surface, and the normalized streamwise velocity *u*. For all turbulence models the flow separates immediately at the leading edge, showing a large separation bubble. No re-attachment takes place. The standard SST *k*-ω model predicts a larger separation bubble, the region of retarded flow is wider and more extended downstream. Modelling transition with the additional *γ*-Re_θ_ model clearly influences the results, and the result are sensitive to the inflow conditions, confirming the difficulty of modelling flow conditions at this Reynolds number regime (*37*).

Based on these CFD results, we estimated the lift coefficient, drag coefficient and lift-to-drag ratio for all turbulence model configurations, and compared them with the empirical estimate of the real butterfly (Fig. S4). This shows that all models overestimated the forces compared to the empirical estimate. The cause of this difference can be due to many aspects, including measurement errors in the empirical analysis (e.g. angle-of-attack, dihedral angle) and overestimations of the glide performance by the CFD solver. Regardless, the force estimates are similar for all models, despite the clear differences in flow dynamics (Fig. S3). This suggests that aerodynamic performance estimates are less sensitive to model selection than detailed airflow dynamics. In our comparison, the SST *k*-ω turbulence model (*40*) had the best match with the empirical estimate for all three metrics (*C_L_, C_D_* and *L/D*), and thus we chose this model for all our further analyses.

#### Studying the effect of wing shape on aerodynamics performance in gliding *Morpho* butterflies

Because CFD simulations rely on large computational resources, we restricted the analysis to three *Morpho* species with strongly divergent wing shapes: *M. cisseis*, a canopy species with large triangular wings, *M. rhetenor*, a canopy species with markedly elongated wings, and *M. deidamia*, an understory species with rounded wings.

For *M. cisseis* and *M. deidamia* we determined wing shape during gliding flight based on videography data of our semi-field study. Herefore, we selected the stereoscopic video where a significant glide occurred just under the top-view camera. From the top-view video, we traced the outline of the wing during gliding, which was used to build the virtual wing model for that species (Fig. S2). No such gliding sequences were however available for individuals *M. rhetenor*, and thus for this species we estimated the gliding wing shape using the same overlap between fore- and hindwings observed in the other species when gliding (Fig. S2).

Because we aimed to study the effect of wing planform shape on aerodynamics, we kept all other metrics as similar as possible between simulations. Therefore, for all butterflies, we modelled fluid dynamics around the left wing, and the interaction effect with the right wing was modelled using a symmetry plane (Fig S2D). All wings consisted of a single rigid flat plate of 1 mm thick, with the edges rounded as a semi-circle of 0.5 mm radius (Fig. S2D). This wing of 1 mm thickness is similar to that of the largest veins located on the leading edge, which is a region that is particularly important regarding airflow development and separation (*33, 41*). We ignored wing flexibility effects, the effect of potential gaps between the fore and hind wing, and the effect of the body, as this was not related to our research question.

To remove the effect of Reynolds number on flow dynamics and glide performance, we performed all simulations at a mean-chord-based Reynolds number of Re = *U*_∞_ *c*/ν ∼ 5200, where *U*_∞_ is inflow velocity, *c* is mean wing chord and ν is kinematic viscosity of air (ν = 1.4776 10^-5^ m^2^.s^-1^). Because the three studied species have different wing chord lengths, we achieved this by varying the inflow velocity between species (as *U*_∞_ = 5200 ν/*c*). This resulted in *U*_∞_=1.59 m.s^-1^; for *M. cisseis*, *U*_∞_=1.97 m.s^-1^ for *M. deidamia*, and *U*_∞_=1.98 m.s^-1^ for *M. rhetenor*. All the simulation setups were run using five meshes of increasing resolution, to estimate the numerical uncertainty brought by the spatial discretization.

For each species, we performed gliding flight simulations at angles-of-attacks ranging from 2° to 20°, with a step size of 2°. At each angle-of-attack, we quantified the airflow dynamics and determined the aerodynamic force vector on the wing **F**_aero_. We visualized the airflow dynamics as using the distributions of normalized air pressure *C_p_* and limited streamlines on the top and bottom surfaces of the wing, and as iso-surfaces of constant λ_2_ criterion around the wing. We color-coded these iso-surfaces with the relative eddy viscosity ν*_t_* /ν, where turbulence is modelled for values higher than one (Figs S5-S7).

From the aerodynamic force vector, we determined the lift coefficient, drag coefficient and lift-to-drag ratio as explained above. The uncertainty interval for the aerodynamic forces and corresponding coefficients estimated in our CFD were determined using on a grid refinement exercise (*36*). These uncertainties are depicted as error bars in Figures 4B and S4.

#### Aerodynamic flight performance of gliding *Morpho* butterflies

For each studied butterfly, we determined the lift-to-drag ratio to angle-of-attack curve, which we used to determine the maximum lift-to-drag ratio. For all species, the lift-to-drag ratio was maximum at an angle-of-attack of 6° (Fig 4B), but it was significantly higher for the two canopy species *M. cisseis* (*L*/*D*max = 5.83 (5.78-5.88)), mean (uncertainty interval)) and *M. rhetenor* (*L*/*D*max = 5.71 (5.66-5.76)) than for the understory species *M. deidamia* (*L*/*D*max = 5.18 (5.14-5.22)).

The airflow visualizations (Fig. S5-S7) show why for all species the lift-to-drag ratio was maximum at α*_L/D_*_max_ = 6°. At the angles-of-attack of both 4° and 6°, the airflow separates from the wings immediately at the leading edge, and reattaches before reaching the trailing edge. As a result, all wings produce an attached separation bubble on top of the wing, generally called a Leading Edge Vortex (LEV) (*41*). This LEV increases in size with angle-of-attack until it almost reaches the trailing edge at α_L/Dmax_ = 6°. At angles-of-attack larger αL/D_max_, the LEV grows to beyond the trailing edge, and full airflow separation occurs. The lack of reattachment translates into an increased drag, which explains why *L/D* quickly drops at angles-of-attack beyond 6° (Fig. 4B).

In order to investigate why the maximum lift-to-drag ratio *L*/*D*_max_ is higher for the canopy species, we used aerodynamic theory to study how drag varied with wing shape. The drag of a wing is defined as *C_D_* = *C_D0_* + *C_Di_*, where *C_D0_* is profile drag and *C_Di_* is lift-induced drag. The lift-induced drag predominantly varies with wing shape as it is defined as

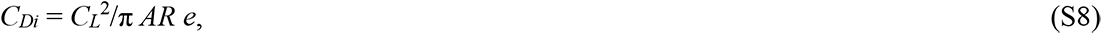

where *AR* and *e* are the aspect-ratio and span efficiency of the wing, respectively. Thus, a wing with a high aspect ratio and a high span efficiency has low lift-induced drag, and consequently a high lift-to-drag ratio. We therefore determined for each wing shape, the aspect ratio based on the wing planforms, and span efficiency based on the CFD data as

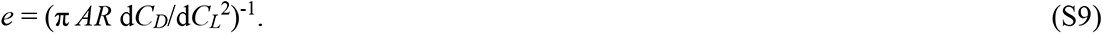

Here, we calculated the derivative d*C_Di_*/d*C_L_*^2^ using a linear fit on the pre-stall *C_D_*-*C_L_*^2^ polar data. Span efficiency is maximum (equal to 1) for an elliptic spanwise lift distribution; any deviation from this elliptic distribution, reduces span efficiency. Therefore, we also determined the spanwise lift distribution of each wing at its maximum lift-to-drag ratio, by integrating the pressure difference across the wing surface along the chord.

The results of this analysis show that the canopy species have a higher *L*/*D*_max_ than the understory species due to lower lift-induced drag. But this reduction in lift-induced drag is achieved by the two canopy species using two distinct aerodynamic mechanisms: *M. cisseis* has a reduced lift-induced drag due to an increased span efficiency, whereas *M. rhetenor* has an increased aspect ratio wing. The airflow dynamics around the wings highlight the mechanisms that cause the increase in span efficiency in *M. cisseis* (Figs S5-S7). At α*L/D*_max_, all three *Morpho* species produce distinct wing tip vortices, but *M. rhetenor* and *M. deidamia* both also produce streamwise-oriented vortices at the location where there is a steep spanwise variation in wing chord length. For *M. rhetenor* this is at approximately two-third wing span, where the elongated wing section with low chord length starts (Fig. S6). For *M. deidamia*, this is at the wing root section, where the wing chord steeply decreases (Fig. S7). In contrast, *M. cisseis* does not produce such streamwise-oriented vortices along the span, except for the tip vortices, as it does not have such rapid spanwise variation in wing chord length (Fig. S5). This explains the relatively small spanwise variation in spanwise lift distribution and the resulting high span efficiency (Fig. 4).

## Supplementary Figures

**Fig. S1.**
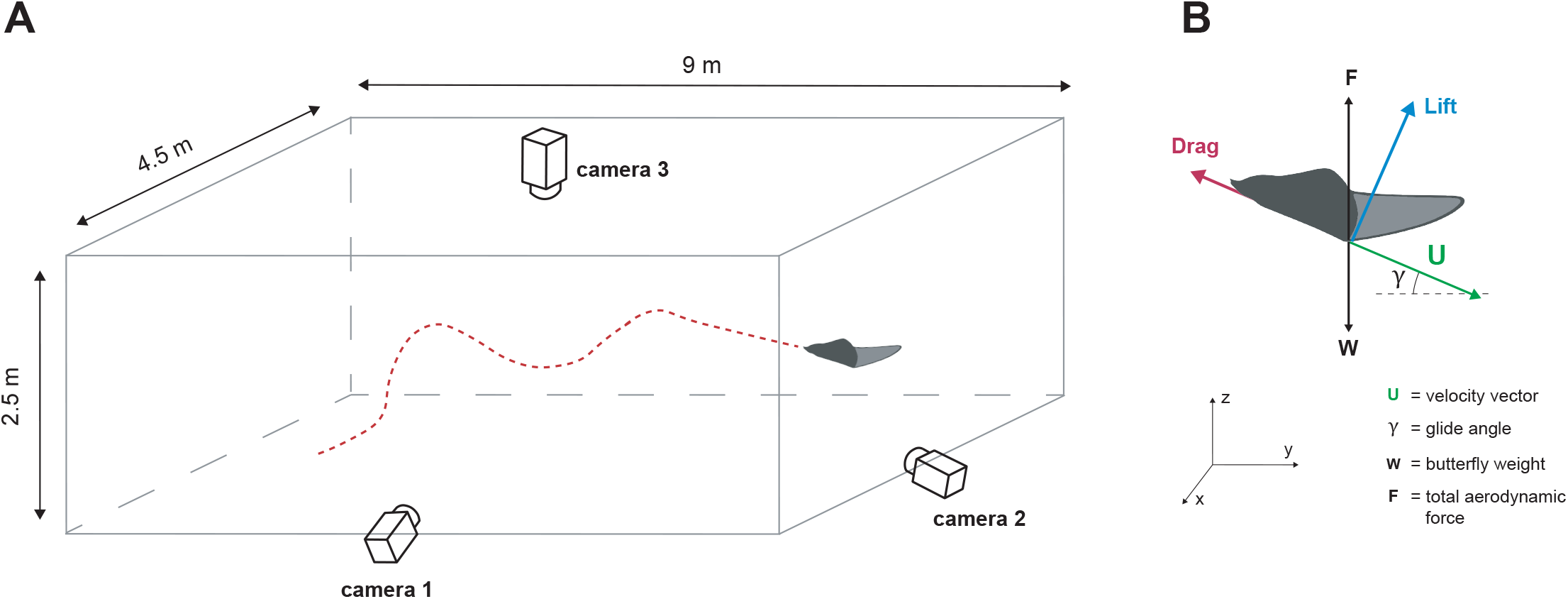
Schematic of the semi-field experimental set-up used to reconstruct the three-dimensional trajectories of butterflies. (**A**) Butterflies were filmed in a large outdoor insectary (9m × 4m × 2.5m) made of fine mesh, using three orthogonally positioned video cameras. (**B**) Butterfly flight dynamics was recontructed from the three-dimensional positions throughout the trajectory. Velocity and force vectors are shown during gliding flight.

**Fig. S2.**
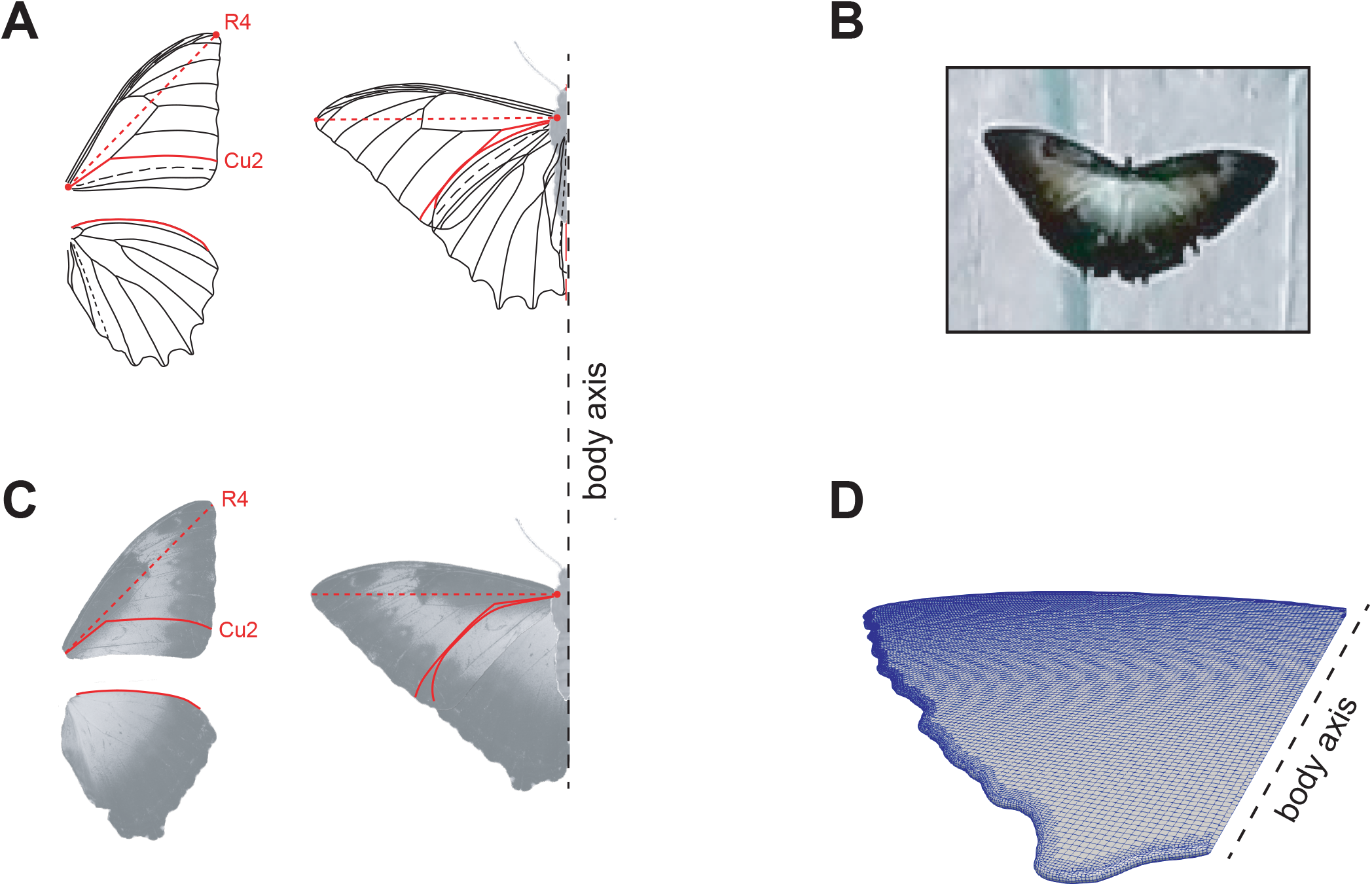
Determining the wing shape of the gliding *Morpho* butterflies for our CFD study. (**A**) The typical vein architecture of *Morpho* butterfly wings (adapted from Blandin 1988, Ref *42*) was use as landmarks to standardize the relative position of the fore- and hindwing during gliding flight. The axis linking the wing base to the vein junction R4 (red dotted line) was positioned perpendicular to the body axis. The front margin of the hindwing was then placed against the Cu2 vein of the forewing (red lines). (**B**) The overlap between forewing and hindwing in (**A**) was determined based on the overlap during gliding flight, estimated based on the top view camera in the semi-field setup. (**C)** Cropped images of the wings from the filmed specimens were used to build the realistic gliding wing shape. (**D**) Three-dimensional model of the gliding wing, including the surface mesh used in the CFD simulations.

**Fig. S3.**
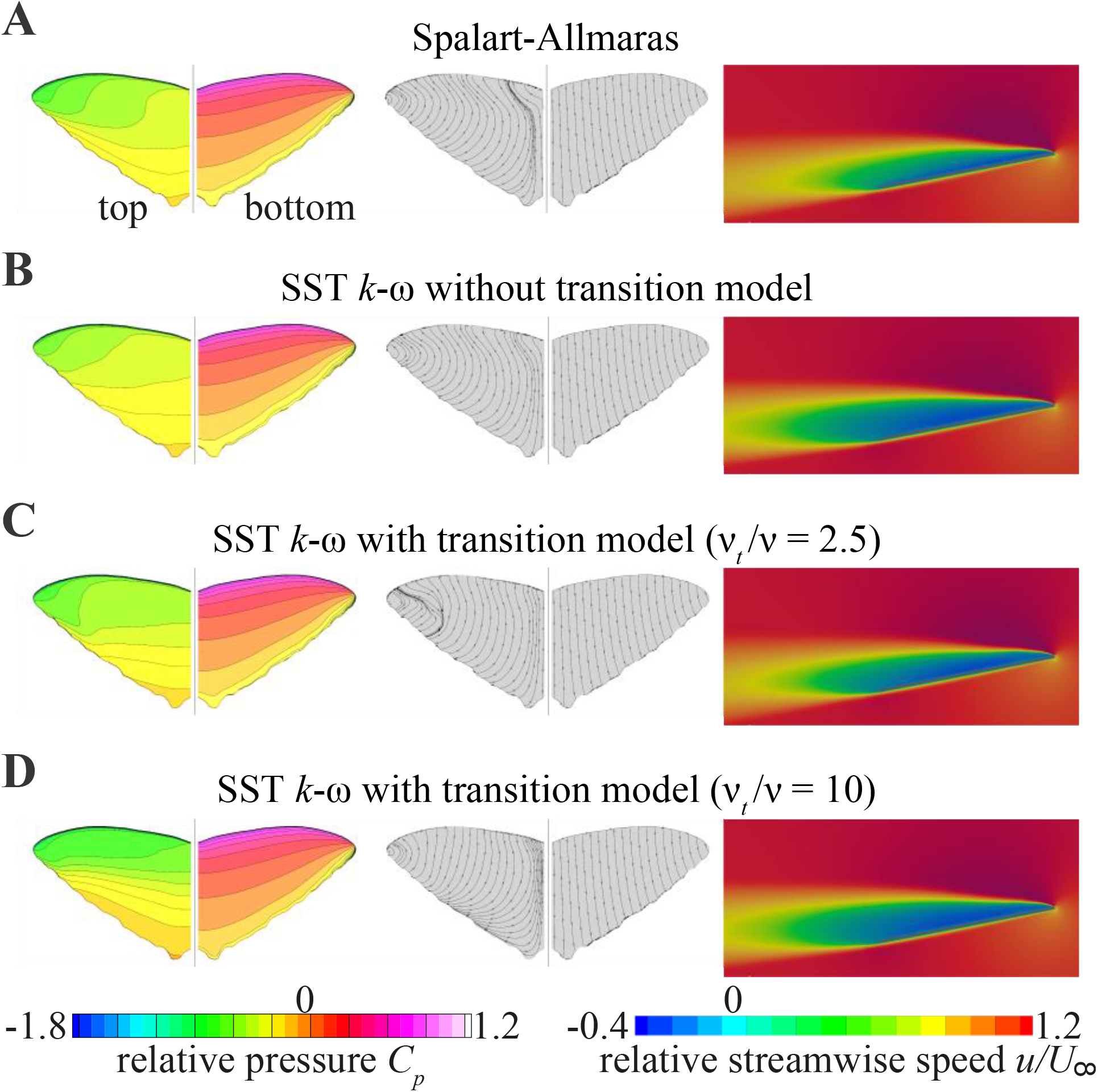
Airflow dynamics around the *M. cisseis* model wing operating at α = 11°, determined using the four turbulence model configurations for our model selection analysis. (**A**-**D**) results of the Spalart-Allmaras model (**A**), SST k-ω without transition model (**B**), SST k-ω with *γ*-Reθ transition model and *ν_t_* = 2.5 inflow condition (**C**), and SST k-ω with *γ*-Reθ transition model and *ν_t_* = 10.0 inflow condition (**D**). Each panel (**A** to **D**) shows the relative air pressure around on the wing surfaces (left), the limiting streamlines on the wing surfaces (middle) and the relative streamwise airspeed around the wing at 1 cm from the wing root (right). For each wing pair figure, the left wing shows the air pressure and streamlines on the dorsal (suction) side, and the right wing shows the air pressure and streamlines on the ventral (pressure) side.

**Fig. S4.**
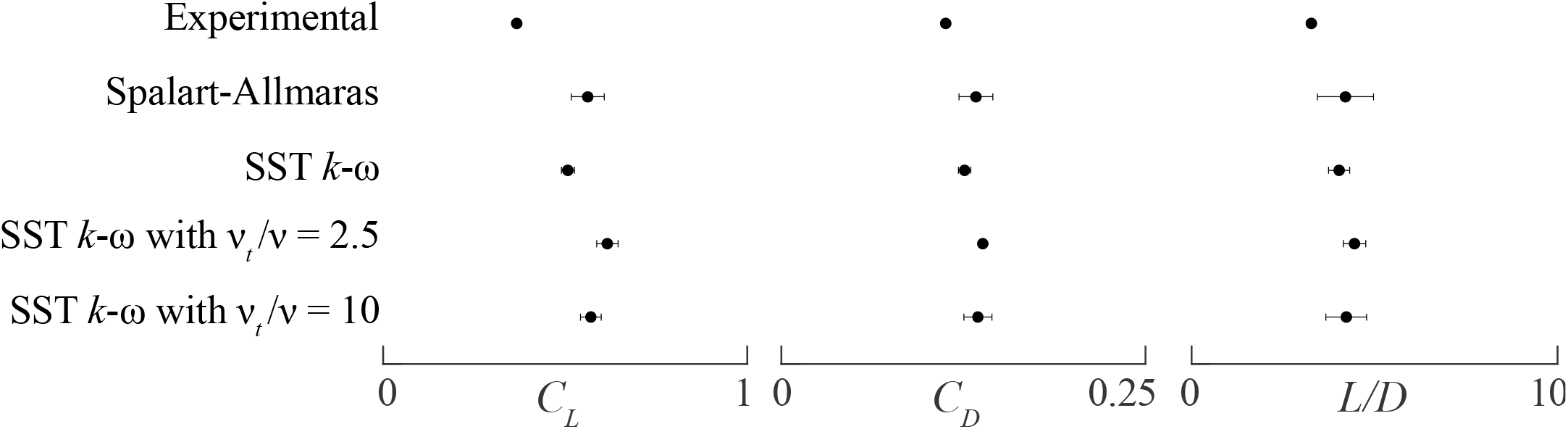
Aerodynamic forces produced by a gliding *M. cisseis*, estimated empirically based on experimental videography data, and estimated numerically using CFD with the various turbulence model configurations. The CFD results are given as mean and uncertainty interval (error bars), estimated using a grid refinement exercise.

**Fig. S5.**
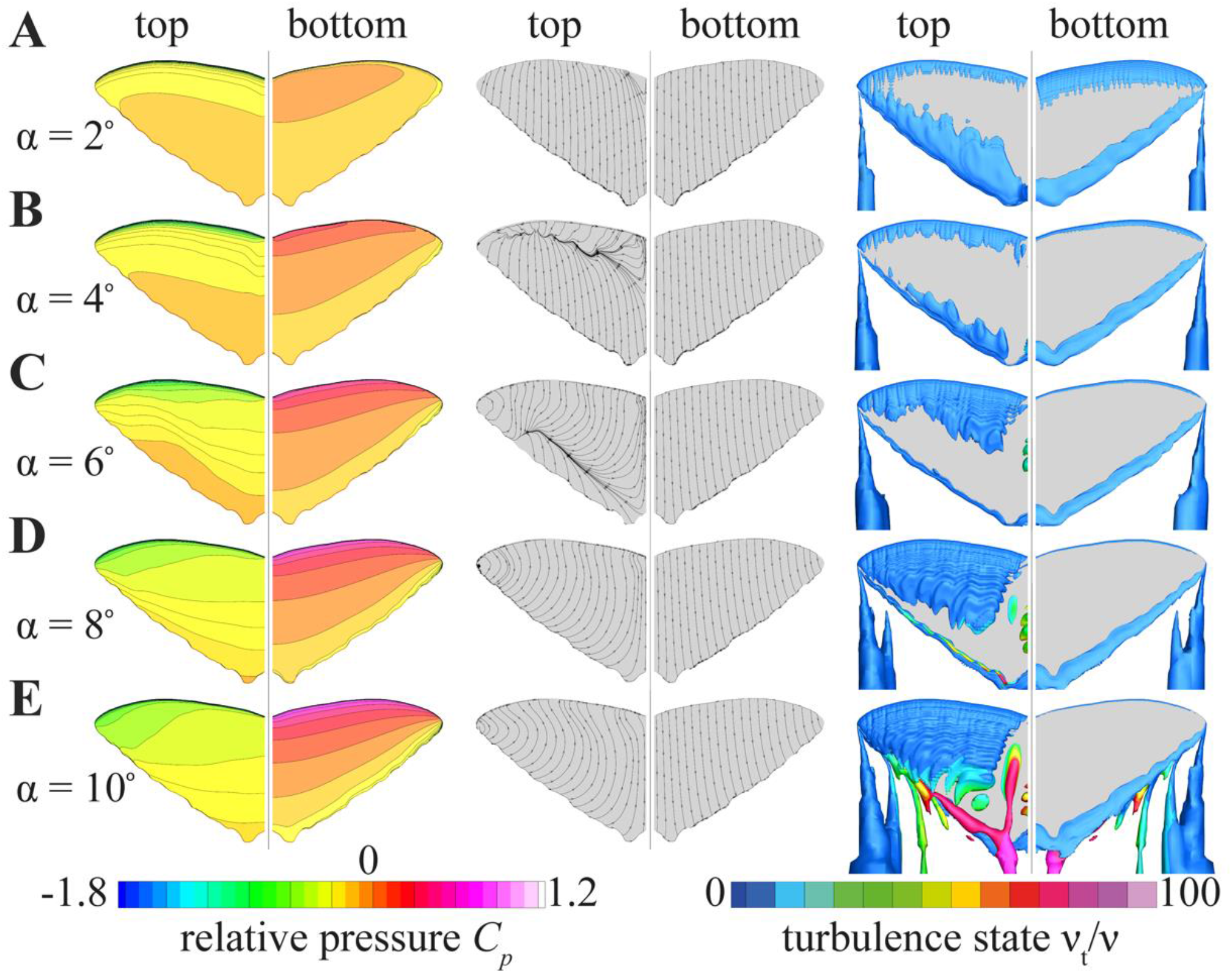
Airflow dynamics around the *M. cisseis* wing gliding at angles-of-attack ranging from 2° to 10° (**A** to **E**, respectively). Each panel shows the relative air pressure *C_p_* around the wing (left), the limiting streamlines (middle), and the vorticity field (right). For each wing pair, the left wing shows the air pressure and streamlines on the dorsal (top) side, and the right wing shows the streamlines on the ventral (bottom) side. The relative air pressure *C_p_* shows the pressure load on the wing, normalized by the free stream pressure. The vortex structures are shown as iso-surfaces of the λ_2_ criterion equal to 0.1. These iso-surfaces are color-coded by relative eddy viscosity ν_*t*_ /ν, where turbulence is modelled for values higher than one.

**Fig. S6.**
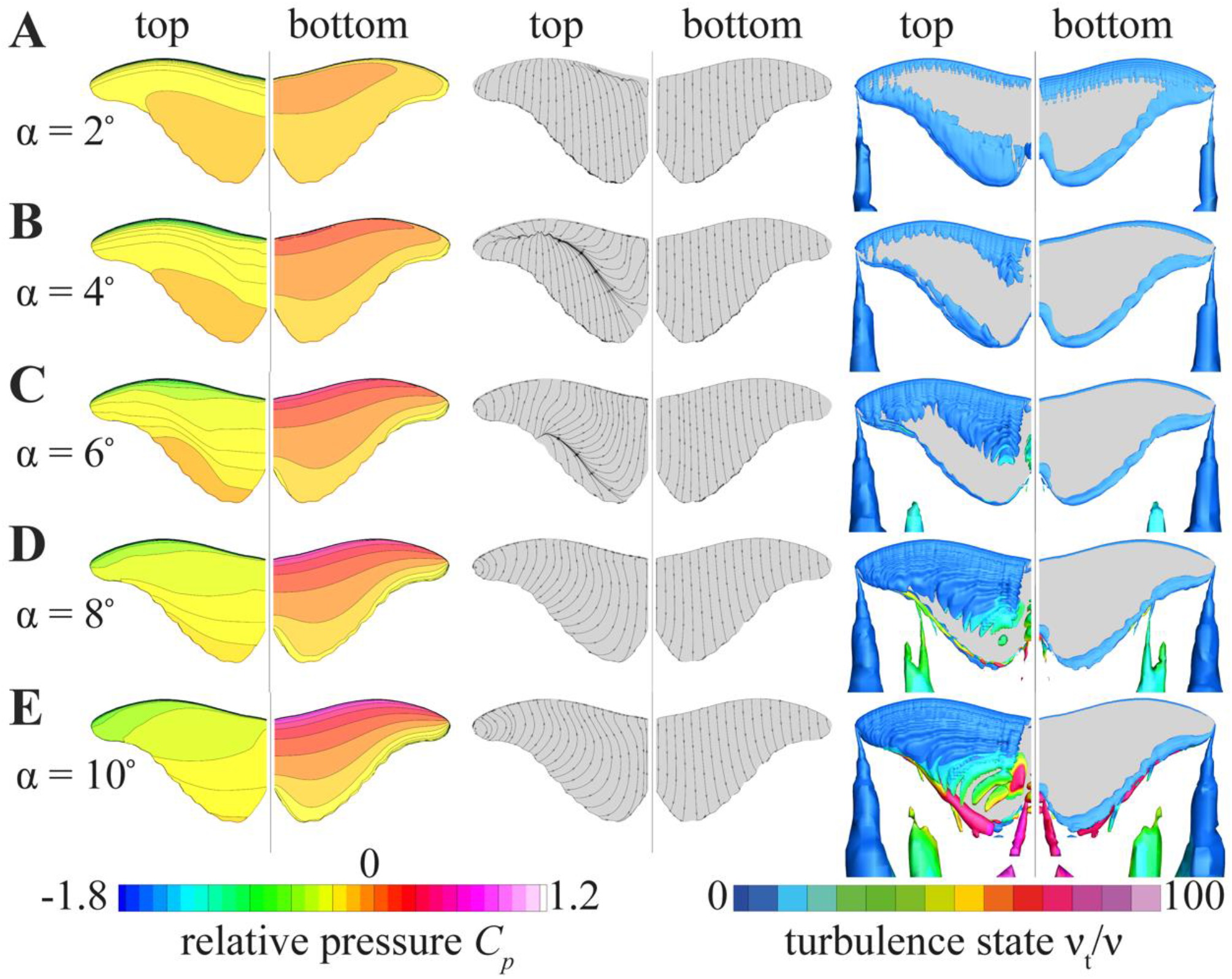
Airflow dynamics around the *M. rhetenor* wing gliding at angles-of-attack ranging from 2° to 10° (**A** to **E**, respectively). Each panel shows the relative air pressure *C_p_* around the wing (left), the limiting streamlines (middle), and the vorticity field (right). For each wing pair, the left wing shows the air pressure and streamlines on the dorsal (top) side, and the right wing shows the streamlines on the ventral (bottom) side. The relative air pressure *C_p_* shows the pressure load on the wing, normalized by the free stream pressure. The vortex structures are shown as iso-surfaces of the λ_2_ criterion equal to 0.1. These iso-surfaces are color-coded by relative eddy viscosity ν_*t*_ /ν, where turbulence is modelled for values higher than one.

**Fig. S7.**
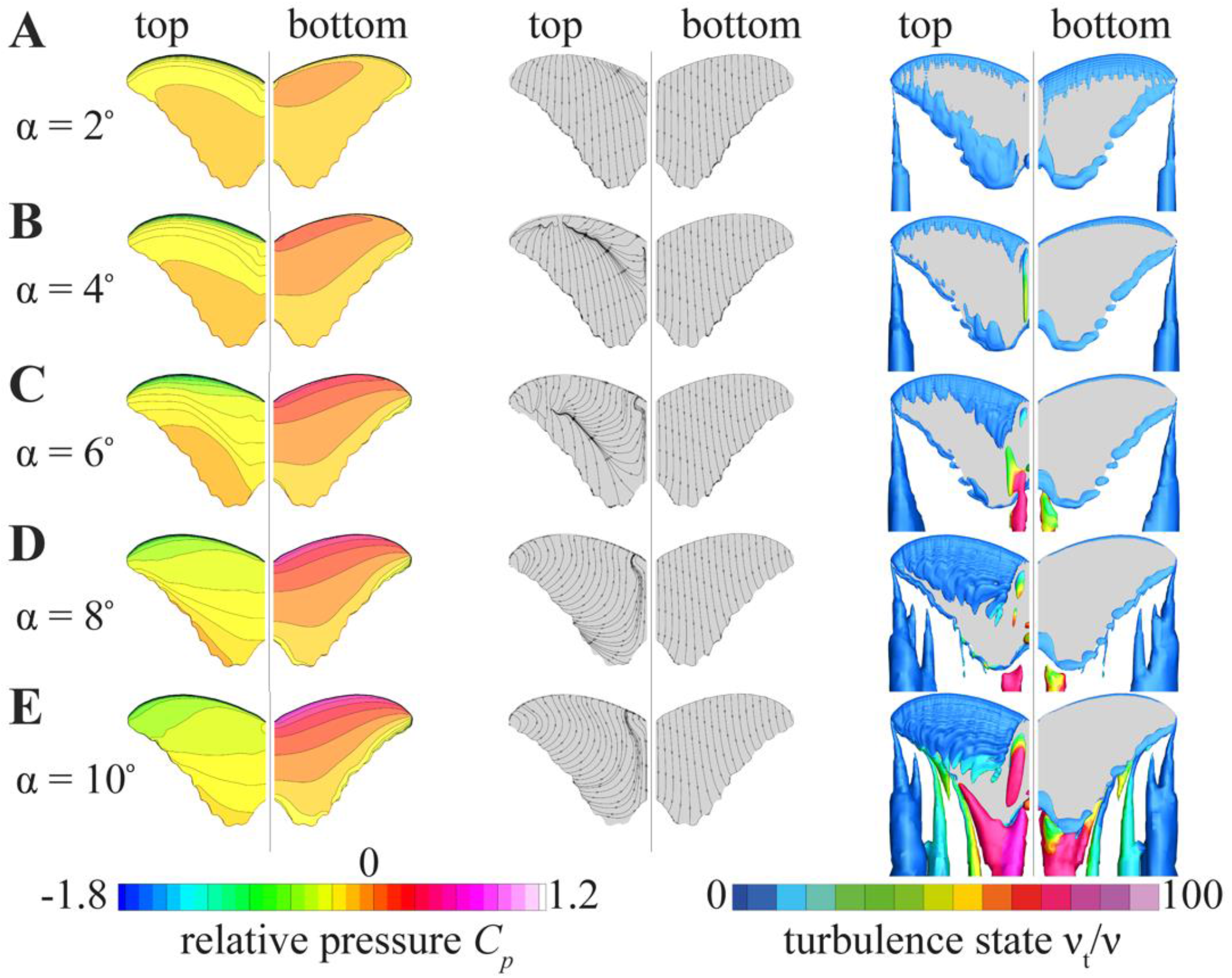
Airflow dynamics around the *M. deidamia* wing gliding at angles-of-attack ranging from 2° to 10° (A to E, respectively). Each panel shows the relative air pressure *C_p_* around the wing (left), the limiting streamlines (middle), and the vorticity field (right). For each wing pair, the left wing shows the air pressure and streamlines on the dorsal (top) side, and the right wing shows the streamlines on the ventral (bottom) side. The relative air pressure *C_p_* shows the pressure load on the wing, normalized by the free stream pressure. The vortex structures are shown as iso-surfaces of the λ_2_ criterion equal to 0.1. These iso-surfaces are color-coded by relative eddy viscosity ν_*t*_ /ν, where turbulence is modelled for values higher than one.

## Supplementary Tables

**Table S1.**
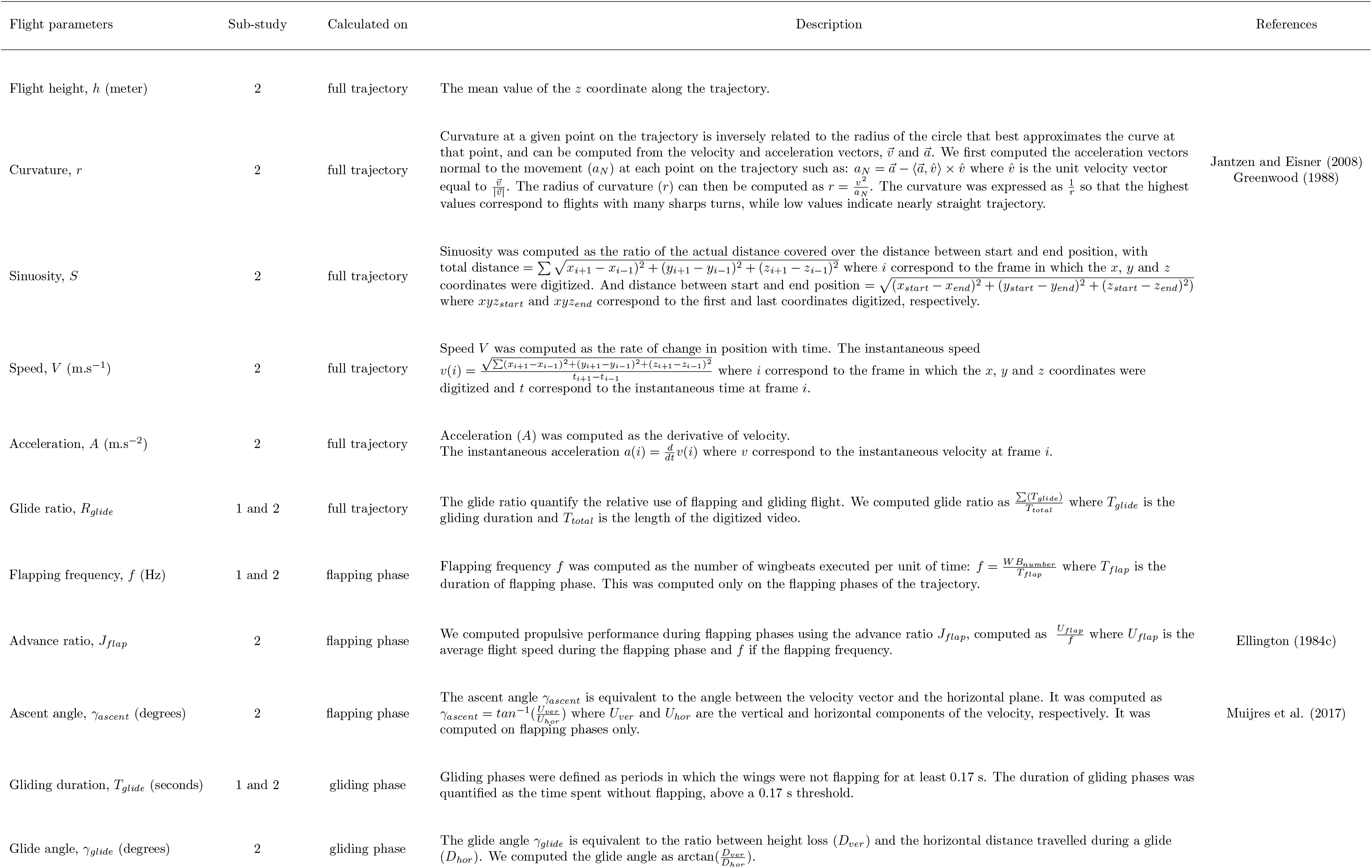
Description of the flight parameters. Parameters were computed in two different sub-studies. Sub-study 1: natural flight sequences of butterflies flying freely in the wild. Sub-study 2: three-dimensional trajectories of butterflies flying in a large outdoor insectary.

**Table S2.**
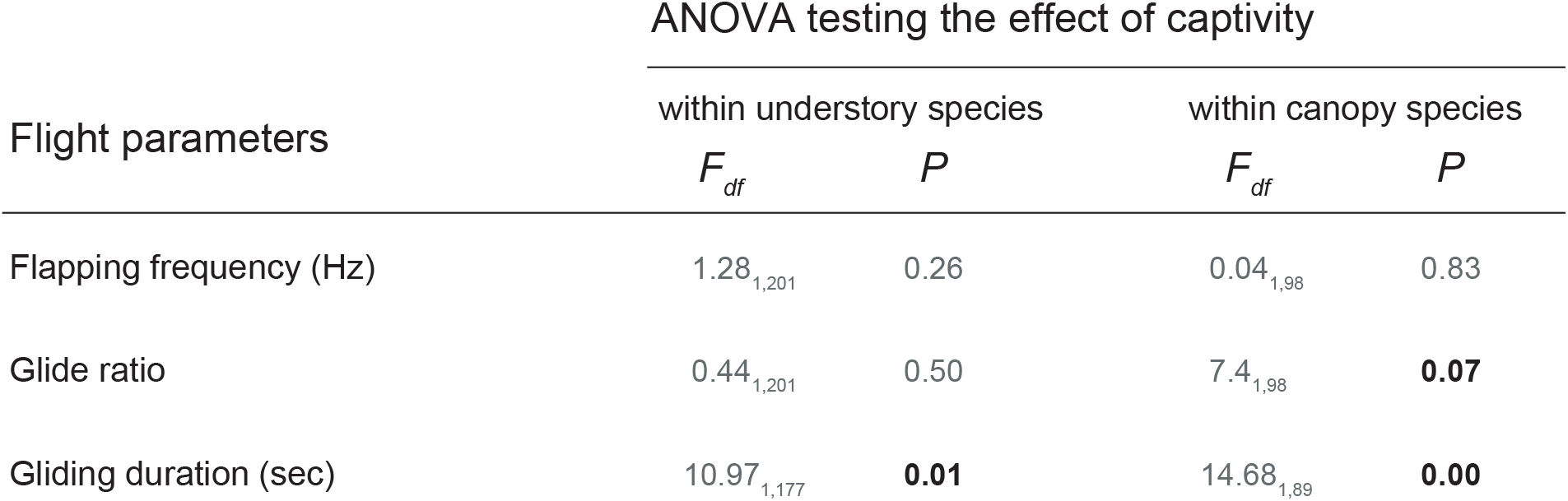
Effect of captivity on the flight parameters measured in both wild and insectary condition tested using ANOVA. The effect of captivity was significant when tested on the set of flight parameters using MANOVA (Wilks’ λ = 0.89; *P* < 0.001).

**Table S3.** Effect of microhabitat on flight behaviour tested using standard ANOVA on individuals and phylogenetic ANOVA on species mean values. Phylogenetic signal computed using Blomberg’s *K* is indicated in the last column. The second column indicates the condition in which flight variables were measured (wild or insectary) and the third column shows the mean value for each flight variable, measured either in canopy or in understory species. The effect of microhabitat on the overall flight behaviour (i.e. on the set of flight parameters) was significant when accounting for the phylogeny (phylogenetic MANOVA: Wilks’ λ = 0.03; *P* = 0.02).

| Flight parameters | Flight condition | Mean value |  | Effect of microhabitat (ANOVA) |  | Effect of microhabitat (phylogenetic ANOVA) |  | Phylogenetic signal |  |
| --- | --- | --- | --- | --- | --- | --- | --- | --- | --- |
| | | canopy | understory | $F_{df}$ | $P$ | $F_{df}$ | $P$ | $K$ | $P$ |
| Flight height (m) | Insectary | 1.42 | 1.28 | 5.3<sub>1,80</sub> | **0.023** | 1.1<sub>1,9</sub> | 0.325 | 0.830 | 0.114 |
| Curvature | Insectary | 0.97 | 0.71 | 6.8<sub>1,80</sub> | **0.010** | 0.3<sub>1,9</sub> | 0.575 | 0.705 | 0.294 |
| Sinuosity | Insectary | 1.25 | 1.19 | 7.9<sub>1,80</sub> | **0.005** | 0.4<sub>1,9</sub> | 0.523 | 1.676 | **0.003** |
| Speed (m.s.<sup>-1</sup>) | Insectary | 1.75 | 2.28 | 16.1<sub>1,80</sub> | **0.0001** | 0.5<sub>1,9</sub> | 0.470 | 1.253 | **0.018** |
| Acceleration (m.s.<sup>-2</sup>) | Insectary | 2.68 | 3.42 | 6.7<sub>1,80</sub> | **0.011** | 0.1<sub>1,9</sub> | 0.671 | 0.929 | 0.09 |
| Glide ratio | Insectary | 0.15 | 0.18 | 0.6<sub>1,80</sub> | 0.424 | 1.4<sub>1,9</sub> | 0.256 | 0.527 | 0.401 |
| Glide ratio | Wild | 0.35 | 0.22 | 9.8<sub>1,134</sub> | **0.002** | 0.6<sub>1,9</sub> | 0.524 | 1.183 | 0.117 |
| Flapping frequency (Hz) | Insectary | 6.14 | 6.03 | 0.2<sub>1,80</sub> | 0.626 | 0.1<sub>1,9</sub> | 0.774 | 0.733 | 0.207 |
| Flapping frequency (Hz) | Wild | 6.23 | 6.20 | 0.0<sub>1,134</sub> | 0.881 | 0.2<sub>1,9</sub> | 0.737 | 0.921 | 0.622 |
| Advance ratio | Insectary | 0.29 | 0.38 | 26.4<sub>1,80</sub> | **0.000001** | 1.0<sub>1,9</sub> | 0.326 | 1.145 | **0.038** |
| Ascent angle (degree) | Insectary | 21.62 | 13.24 | 9.2<sub>1,80</sub> | **0.003** | 6.8<sub>1,9</sub> | **0.021** | 0.953 | **0.006** |
| Gliding duration (sec) | Insectary | 0.24 | 0.25 | 0.0<sub>1,80</sub> | 0.867 | 0.0<sub>1,9</sub> | 0.906 | 0.463 | 0.522 |
| Gliding duration (sec) | Wild | 0.50 | 0.32 | 9.6<sub>1,100</sub> | **0.002** | 1.1<sub>1,9</sub> | 0.466 | 1.231 | **0.042** |
| Glide angle (degree) | Insectary | 6.77 | 10.57 | 10.5<sub>1,80</sub> | **0.001** | 7.6<sub>1,9</sub> | **0.020** | 1.325 | **0.006** |

**Table S4.** Effect of wing loading and forewing aspect-ratio on flight parameters. Significant *p*-value are indicated in bold.

| Flight variables | Wing loading |  |  |  |  |  | Forewing aspect-ratio |  |  |  |  |  |
| --- | --- | --- | --- | --- | --- | --- | --- | --- | --- | --- | --- | --- |
|  | Simple regression |  |  | Phylogenetic regression |  |  | Simple regression |  |  | Phylogenetic regression |  |  |
|  | *R* | *F*<sub>df</sub> | *P* | *R* | *F*<sub>df</sub> | *P* | *R* | *F*<sub>df</sub> | *P* | *R* | *F*<sub>df</sub> | *P* |
| Flight height | -0.15 | 1.9<sub>1,80</sub> | 0.169 | -0.01 | 0.0<sub>1,9</sub> | 0.876 | 0.01 | 0.3<sub>1,80</sub> | 0.574 | 0.00 | 0.4<sub>1,9</sub> | 0.523 |
| Curvature | -0.30 | 8.0<sub>1,80</sub> | **0.005** | -0.31 | 5.5<sub>1,9</sub> | **0.042** | 0.01 | 0.1<sub>1,80</sub> | 0.734 | 0.12 | 0.0<sub>1,9</sub> | 0.927 |
| Sinuosity | -0.25 | 5.2<sub>1,80</sub> | **0.025** | -0.11 | 0.0<sub>1,9</sub> | 0.916 | 0.02 | 0.6<sub>1,80</sub> | 0.440 | 0.11 | 0.0<sub>1,9</sub> | 0.901 |
| Speed | 0.42 | 17.4<sub>1,80</sub> | **0.000** | 0.61 | 16.9<sub>1,9</sub> | **0.002** | -0.17 | 2.4<sub>1,80</sub> | 0.124 | 0.12 | 0.0<sub>1,9</sub> | 0.978 |
| Acceleration | 0.30 | 7.8<sub>1,80</sub> | **0.006** | 0.78 | 36.2<sub>1,9</sub> | **0.000** | -0.14 | 1.6<sub>1,80</sub> | 0.202 | 0.03 | 0.2<sub>1,9</sub> | 0.674 |
| Glide ratio | -0.16 | 2.2<sub>1,80</sub> | 0.143 | 0.01 | 0.4<sub>1,9</sub> | 0.515 | -0.24 | 5.1<sub>1,80</sub> | **0.026** | -0.22 | 3.9<sub>1,9</sub> | 0.079 |
| Flapping frequency | 0.47 | 23.3<sub>1,80</sub> | **0.000** | 0.02 | 0.9<sub>1,9</sub> | 0.350 | 0.28 | 7.1<sub>1,80</sub> | **0.009** | 0.23 | 4.1<sub>1,9</sub> | 0.073 |
| Advance ratio | 0.22 | 4.1<sub>1,80</sub> | **0.046** | 0.30 | 5.2<sub>1,9</sub> | **0.048** | -0.29 | 7.5<sub>1,80</sub> | **0.007** | 0.05 | 0.8<sub>1,9</sub> | 0.384 |
| Ascent angle | 0.10 | 0.9<sub>1,80</sub> | 0.342 | 0.27 | 4.7<sub>1,9</sub> | 0.057 | 0.33 | 9.7<sub>1,80</sub> | **0.002** | 0.23 | 3.9<sub>1,9</sub> | 0.076 |
| Gliding duration | -0.24 | 5.1<sub>1,80</sub> | **0.026** | 0.05 | 0.0<sub>1,9</sub> | 0.936 | -0.23 | 4.5<sub>1,80</sub> | **0.036** | 0.06 | 2.0<sub>1,9</sub> | 0.182 |
| Glide angle | -0.02 | 0.0<sub>1,80</sub> | 0.803 | 0.01 | 0.9<sub>1,9</sub> | 0.360 | -0.35 | 11.0<sub>1,80</sub> | **0.001** | -0.13 | 2.5<sub>1,9</sub> | 0.147 |

**Table S5.** The number of cells of each numerical grid used in our grid refinement exercise and for estimating the numerical / discretization uncertainties (*36*). We used four refinement steps for the model selection analysis, and we used five refinement steps for our final study comparing the glide performance of three *Morpho* species. The simulations at the finest grid (in bold) were used for determining aerodynamic forces and airflow dynamics; the complete set of simulations were used to estimate the uncertainties in aerodynamic force production.

| Grid number | Model selection | Wing shape comparison |  |  |
| --- | --- | --- | --- | --- |
|  |  | *M. cisseis* | *M. deidamia* | *M. rhetenor* |
| 1 | 1,157,340 | 698,804 | 481,503 | 645,452 |
| 2 | 3,785,147 | 1,810,196 | 1,249,733 | 1,662,434 |
| 3 | 8,406,881 | 3,377,716 | 2,312,933 | 3,094,332 |
| 4 | **15,738,960** | 5,715,510 | 3,911,936 | 5,223,167 |
| 5 | - | **9,658,292** | **6,643,215** | **8,854,294** |

**Table S6.** Morphological and flight characteristics of *M. cisseis, M. rheteno* and *M. deidamia* during the gliding flight sequences recorded in free flight in the semi-field setup, and used as input for the CFD study

| Characteristics |  | Species |  |  |
| --- | --- | --- | --- | --- |
|  |  | *M. cisseis* | *M. deidamia* | *M. rhetenor* |
| Mean body mass | [g] | 0.58 | 0.64 | 0.54 |
| Mean wing chord length | [cm] | 4.82 | 3.91 | 3.88 |
| Wing surface area | [cm<sup>2</sup>] | 44.1 | 25.8 | 36.5 |
| Root-to-tip wingspan | [cm] | 8.75 | 7.56 | 9.41 |
| Wing aspect ratio | [ - ] | 3.63 | 3.86 | 4.85 |
| Mean gliding speed | [m.s<sup>-1</sup>] | 1.59 | 1.97 | 1.98 |

**Movie S1**

Stereoscopic flight sequence of an individual *Morpho cisseis* (canopy species) showing typical flap-gliding flight.

**Movie S2**

Stereoscopic flight sequence of an individual *Morpho godartii* (understory species) showing powerful flapping flight.

## References and Notes

1. G. K. Taylor, A. L. Thomas, Dynamic flight stability in the desert locust *Schistocerca gregaria*. Journal of Experimental Biology 206, 2803 (2003).

2. S. Altizer, A. K. Davis, Populations of monarch butterflies with different migratory behaviors show divergence in wing morphology. Evolution: International Journal of Organic Evolution 64, 1018 (2010).

3. T. M. Casey, Flight energetics of sphinx moths: power input during hovering flight. Journal of Experimental Biology 64, 529 (1976).

4. M. E. Dillon, R. Dudley, Allometry of maximum vertical force production during hovering flight of neotropical orchid bees (Apidae: Euglossini). Journal of Experimental Biology 207, 417 (2004).

5. B. A. Holloway, Pollen-feeding in hover-flies (Diptera: Syrphidae). New Zealand journal of zoology 3, 339 (1976).

6. F. T. Muijres, M. J. Elzinga, J. M. Melis, M. H. Dickinson, Flies evade looming targets by executing rapid visually directed banked turns. Science 344, 172 (2014).

7. A. Cribellier et al., Flight behaviour of malaria mosquitoes around odour-baited traps: capture and escape dynamics. Royal Society Open Science 5, 180246 (2018).

8. A. P. Willmott, C. P. Ellington, The mechanics of flight in the hawkmoth *Manduca sexta*. II. Aerodynamic consequences of kinematic and morphological variation. Journal of Experimental Biology 200, 2723 (1997).

9. H. W. Bates, The naturalist on the river Amazon., (John Murray, London, 1864).

10. F. T. Muijres, P. Henningsson, M. Stuiver, A. Hedenström, Aerodynamic flight performance in flap-gliding birds and bats. Journal of theoretical biology 306, 120 (2012).

11. P. DeVries, C. M. Penz, R. I. Hill, Vertical distribution, flight behaviour and evolution of wing morphology in Morpho butterflies. Journal of Animal Ecology 79, 1077 (2010).

12. N. Chazot et al., Morpho morphometrics: Shared ancestry and selection drive the evolution of wing size and shape in Morpho butterflies. Evolution 70, 181 (2016).

13. U. M. Norberg, Structure, form, and function of flight in engineering and the living world. Journal of morphology 252, 52 (2002).

14. P. Henningsson, R. J. Bomphrey, Span efficiency in hawkmoths. Journal of the Royal Society Interface 10, 20130099 (2013).

15. F. T. Muijres, G. R. Spedding, Y. Winter, A. Hedenström, Actuator disk model and span efficiency of flapping flight in bats based on time-resolved PIV measurements. Experiments in fluids 51, 511 (2011).

## References

1. P. Blandin, B. Purser, Evolution and diversification of Neotropical butterflies: Insights from the biogeography and phylogeny of the genus Morpho Fabricius, 1807 (Nymphalidae: Morphinae), with a review of the geodynamics of South America. Tropical Lepidoptera Research 23, 62 (2013).

2. M. J. Fernández, M. E. Driver, T. L. Hedrick, Asymmetry costs: effects of wing damage on hovering flight performance in the hawkmoth Manduca sexta. Journal of Experimental Biology, jeb. 153494 (2017).

3. C. Le Roy, R. Cornette, V. Llaurens, V. Debat, Effects of natural wing damage on flight performance in Morpho butterflies: what can it tell us about wing shape evolution? Journal of Experimental Biology 222, jeb204057 (2019).

4. R. Hartley, A. Zisserman, Multiple view geometry in computer vision. (Cambridge university press, 2003).

5. T. L. Hedrick, Software techniques for two-and three-dimensional kinematic measurements of biological and biomimetic systems. Bioinspiration & biomimetics 3, 034001 (2008).

6. D. H. Theriault et al., A protocol and calibration method for accurate multi-camera field videography. Journal of Experimental Biology, jeb. 100529 (2014).

7. F. T. Muijres, M. J. Elzinga, J. M. Melis, M. H. Dickinson, Flies evade looming targets by executing rapid visually directed banked turns. Science 344, 172 (2014).

8. . D. T. Greenwood, Principles of dynamics. (Prentice-Hall Englewood Cliffs, NJ, 1988).

9. C. Ellington, The aerodynamics of hovering insect flight. III. Kinematics. Philosophical Transactions of the Royal Society of London B: Biological Sciences 305, 41 (1984).

10. F. L. Bookstein, Morphometric tools for landmark data: geometry and biology. (Cambridge University Press, 1997).

11. D. C. Adams, F. J. Rohlf, D. E. Slice, Geometric morphometrics: ten years of progress following the ‘revolution’. Italian Journal of Zoology 71, 5 (2004).

12. . C. J. Breuker, M. Gibbs, S. Van Dongen, T. Merckx, H. Van Dyck, in Morphometrics for Nonmorphometricians. (Springer, 2010), pp. 271–287.

13. D. Outomuro, D. C. Adams, F. Johansson, Wing shape allometry and aerodynamics in calopterygid damselflies: a comparative approach. BMC evolutionary biology 13, 118 (2013).

14. N. Chazot et al., Morpho morphometrics: Shared ancestry and selection drive the evolution of wing size and shape in Morpho butterflies. Evolution 70, 181 (2016).

15. A. Fraimout et al., Phenotypic plasticity of Drosophila suzukii wing to developmental temperature: implications for flight. Journal of Experimental Biology 221, jeb166868 (2018).

16. P. Gunz, P. Mitteroecker, Semilandmarks: a method for quantifying curves and surfaces. Hystrix, the Italian Journal of Mammalogy 24, 103 (2013).

17. F. J. Rohlf, The tps series of software. Hystrix 26, 1 (2015).

18. F. J. Rohlf, D. Slice, Extensions of the Procrustes method for the optimal superimposition of landmarks. Systematic Biology 39, 40 (1990).

19. D. C. Adams, E. Otárola-Castillo, geomorph: an R package for the collection and analysis of geometric morphometric shape data. Methods in Ecology and Evolution 4, 393 (2013).

20. R. B. Srygley, R. Dudley, Correlations of the position of center of body mass with butterfly escape tactics. Journal of Experimental Biology 174, 155 (1993).

21. K. Berwaerts, H. Van Dyck, P. Aerts, Does flight morphology relate to flight performance? An experimental test with the butterfly Pararge aegeria. Functional ecology 16, 484 (2002).

22. V. Bonhomme, S. Picq, C. Gaucherel, J. Claude, Momocs: outline analysis using R. Journal of Statistical Software 56, 1 (2014).

23. S. P. Blomberg, T. Garland Jr, A. R. Ives, Testing for phylogenetic signal in comparative data: behavioral traits are more labile. Evolution 57, 717 (2003).

24. D. C. Adams, A generalized K statistic for estimating phylogenetic signal from shape and other high-dimensional multivariate data. Systematic Biology 63, 685 (2014).

25. R. P. Freckleton, P. H. Harvey, M. Pagel, Phylogenetic analysis and comparative data: a test and review of evidence. The American Naturalist 160, 712 (2002).

26. L. J. Harmon, J. T. Weir, C. D. Brock, R. E. Glor, W. Challenger, GEIGER: investigating evolutionary radiations. Bioinformatics 24, 129 (2008).

27. D. C. Adams, R. N. Felice, Assessing trait covariation and morphological integration on phylogenies using evolutionary covariance matrices. PloS one 9, e94335 (2014).

28. F. J. Rohlf, M. Corti, Use of two-block partial least-squares to study covariation in shape. Systematic Biology 49, 740 (2000).

29. L. R. Monteiro, Multivariate regression models and geometric morphometrics: the search for causal factors in the analysis of shape. Systematic Biology 48, 192 (1999).

30. C. P. Klingenberg, Size, shape, and form: concepts of allometry in geometric morphometrics. Development genes and evolution 226, 113 (2016).

31. H. H. Hu, in Fluid mechanics. (Elsevier, 2012), pp. 421-472.

32. H. Liu, K. Kawachi, A numerical study of insect flight. Journal of Computational Physics 146, 124 (1998).

33. S. P. Sane, The aerodynamics of insect flight. Journal of Experimental Biology 206, 4191 (2003).

34. J. Young, S. M. Walker, R. J. Bomphrey, G. K. Taylor, A. L. Thomas, Details of insect wing design and deformation enhance aerodynamic function and flight efficiency. Science 325, 1549 (2009).

35. L. Eça, C. M. Klaij, G. Vaz, M. Hoekstra, F. S. Pereira, On code verification of RANS solvers. Journal of Computational Physics 310, 418 (2016).

36. L. Eça, M. Hoekstra, A procedure for the estimation of the numerical uncertainty of CFD calculations based on grid refinement studies. Journal of Computational Physics 262, 104 (2014).

37. G. Spedding, A. Hedenström, J. McArthur, M. Rosén, The implications of low-speed fixed-wing aerofoil measurements on the analysis and performance of flapping bird wings. Journal of Experimental Biology 211, 215 (2008).

38. F. R. Menter, M. Kuntz, R. Langtry, Ten years of industrial experience with the SST turbulence model. Turbulence, heat and mass transfer 4, 625 (2003).

39. R. B. Langtry, et al., A correlation-based transition model using local variables—Part II: Test cases and industrial applications. (2006).

40. F. Menter, in 23rd fluid dynamics, plasmadynamics, and lasers conference. (1993), pp. 2906.

41. C. P. Ellington, C. Van Den Berg, A. P. Willmott, A. L. Thomas, Leading-edge vortices in insect flight. Nature 384, 626 (1996).

42. P. Blandin, The Genus Morpho (Lepidoptera, Nymphalidae). (Sciences Nat, 1988).

